# DNA Nanostar Structures with Tunable Auxetic Properties

**DOI:** 10.1101/2023.12.22.573109

**Authors:** Yancheng Du, Ruixin Li, Anirudh S. Madhvacharyula, Alexander A. Swett, Jong Hyun Choi

**Affiliations:** School of Mechanical Engineering, Purdue University, West Lafayette, Indiana 47907, United States

**Keywords:** Auxetic, Metamaterials, DNA nanotechnology, DNA origami, DNA self-assembly

## Abstract

Auxetic structures are unique with a negative Poisson’s ratio. Unlike regular materials, they response to external loading with simultaneous expansion or compression in all directions, rendering powerful properties advantageous in diverse applications from manufacturing to space engineering. The auxetic behaviors are determined by structural design and architecture. Such structures have been discovered in natural crystals and demonstrated synthetically with bulk materials. Recent development of DNA-based structures has pushed the unit cell size to nanometer scale. DNA nanotechnology utilizes sequence complementarity between nucleotides. By combining sequence designs with programmable self-assembly, it is possible to construct complex structures with nanoscale accuracy and to perform dynamic reconfigurations. Herein, we report a novel design of auxetic nanostars with sliding behaviors using DNA origami. Our proposed structure, inspired by an Islamic pattern, demonstrates a unit cell with two distinct reconfigurations by programming directed sliding mechanisms. Compared to previous metamaterials, the DNA nanostars show an architecture with tunable auxetic properties for the first time. We envision that this strategy may form the basis of novel metastructures with adaptability and open new possibilities in bioengineering.

## INTRODUCTION

Auxetic metamaterials are synthetic architectures that respond to external loadings in a unique manner. Regular materials under compressive (or extensive) forces will expand (or contract) in orthogonal directions. In contrast, auxetic structures deform simultaneously in all directions. Poisson’s ratio measures such a property which describes deformation behaviors quantitatively:

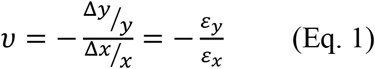

where *ε*_*x*_ and *ε*_*y*_ are strains, and *x* is the loading direction.^1^ Ordinary materials thus have positive Poisson’s ratios, while auxetic structures show negative values. Architected metamaterials have several distinct advantages including light-weight high-strengths^2-4^ and the ability to absorb impact forces^5,6^. They are widely used in design and manufacturing ranging from commodities (e.g., shoes and clothes) to aerospace engineering.^7-9^

Most metamaterials have periodic cellular structures and their auxetic behaviors arise from unit cell designs, which will deform in unison upon external forces. The unit cells range over various length scales. For example, naturally occurring crystals such as α-cristobalite and cubic metals demonstrate auxetic properties with unit cells of sub-nanometer sizes.^10-12^ Conventional auxetics have been manufactured with metals,^13^ polymers,^14^ and other materials^15^ in larger scales from microns to centimeters. There is a lack of studies in auxetic metamaterials at the nanoscale.^16^ Recently, Li *et al*. bridged this gap in lengthscale.^17,18^ In their studies, nanoscale auxetic units were constructed using DNA origami exploiting sequence complementarity of DNA molecules. DNA self-assembly has been developed as a powerful bottom-up strategy with excellent programmability and precision^19^ and demonstrated for complex architectures,^20-24^ reconfigurable designs,^25-28^ and dynamics processes.^29-32^ Their origami architectures were designed with ‘jack’ edges, whose lengths were adjusted by two-step DNA reactions for global structural transformations. The ‘chemical deformation’ resulted in negative Poisson’s ratio (NPR) behaviors.

While the work opened new opportunities for nanoscale metamaterials, the structures were limited in that their auxetic deformations were pre-determined and could not change. In fact, this invariable mechanics is also similar for most metastructures at other lengthscales. Herein we ask if it is possible to program distinct pathways of NPR reconfigurations. To achieve such tunable auxetic properties, we propose a novel design of origami-based DNA nanostars that can reconfigure in different directions upon external loading via chemical deformations. The DNA architecture was inspired by one of the Islamic pattern designs that have been used in arts and buildings. Our DNA origami design consists of 3 nanostars that can slide against each other in two distinct directions, thus resulting in two NPR values. We have investigated their structures and behaviors with coarse-grained molecular dynamics (MD) simulations and atomic force microscopy (AFM). This work opens a new horizon towards smart materials with adaptive mechanical properties for applications involving complex and ever-changing environments.

## EXPERIMENTAL SECTION

### Computer-Aided Origami Design

The design of the wireframe DNA origami was conducted using caDANano2.^33,34^ The edges were designed as four double-stranded (ds) bundles with staples routed around to increase rigidity. The vertices were designed with single-stranded (ss) DNA to allow flexibility in turning angles. In our design, regular right-handed B-form DNA was assumed to have a rise on the axis of 0.332 nm/bp. Positional staples on edges used for performing sliding actions contained binding domains of 10 nt. For strand displacement reactions, 8-nt ss-toeholds were added at one end of the strands. Detailed sequence information is presented in **Table S1 and S2**.

### DNA Origami Synthesis

All DNA origami structures were prepared at a concentration of 5 nM scaffold DNA with 4× staple strands in 1× TAE buffer solution (containing 40 mM trisaminomethane, 20 mM acetic acid, and 1 mM ethylenediaminetetraacetic acid (EDTA) disodium salt with pH ∼ 8). A final concentration of 6 mM Mg^2+^ was provided. The mixture was put in a BIO-RAD S1000 Thermal Cycler under a custom-designed thermal cycle. The sample was heated to 75 °C for 18 minutes and then cooled down at a ramping rate of -0.1 °C/15 mins until it reached 25 °C. The solution was then stored at 4 °C before further experiments.

### Sliding Experiment

Toehold-mediated strand displacement was used for origami sliding motions. The sliding was performed in two steps: (1) removal of previous positional staples and (2) insertion of new positional staples. The first step was carried out by adding 50× releaser strands and incubating the mixture in the thermal cycler at 40 °C for 12 hours before cooling down to 25 °C at a rate of - 0.1°C/6 seconds. Then the mixture was purified by using centrifugal filters (Amicon) to remove excess strands. For centrifugation, approximately 55 μL of DNA origami mixture was added to the filter and additional TAE buffer with 6 mM Mg^2+^ was added to reach a final volume of 500 μL. Then the mixture was centrifuged at 5,000 rpm for 3 minutes and the solution remaining in the filter was retrieved. After the first step, the origami structures were released to an undefined position. In the second step, new staples defining the relative positions of the three-stars were added at a 10-fold higher concentration. The mixture was subsequently reannealed in the thermal cycler at 40 °C for 12 hours and then cooled down to 25 °C at a rate of -0.1°C/6 seconds. The sample was then stored at 4 °C for further measurements or experiments.

### Coarse-grained MD Simulations

We performed MD simulations to evaluate the DNA design and verify structural integrity. In the coarse-grained MD model, pseudo-atoms are used to represent a group of atoms in order to reduce the complexity of calculations. For a predetermined simulation time, particles representing the structures follow designated interactions to demonstrate the development of the system dynamically. In our study, oxDNA platform^35,36^ was used for computing the equilibrium conformations of the origami structures. In the oxDNA model, DNA strands are represented by a string of rigid nucleotides, where multiple interactions are taken into consideration including sugar-phosphate backbone connectivity, excluded volume, hydrogen bonding, etc. To perform the computation, we used the designed static structures from caDNAno and converted them into topology and configuration files using real sequence information as initial conformations in oxDNA. The environmental parameters were set based on the experimental conditions. Temperature was the same as in experiments, while magnesium concentration of [Mg^2+^] = 6 mM was replaced by [Na^+^] = 0.5 M due to limited options in the computational platform. The initial configurations of DNA origami were first relaxed by setting the phosphate backbone connection 10 times stronger to pull the edges to their places. Then, a threshold of 3% relative fluctuations was set to start the second stage of simulations, where the structures were computed for more than 10^6^ steps to reach an equilibrium state for observation. For each conformation, the simulation took about two days. The resulting structures were visualized with oxView^37^ and the quantitative measurements were performed on the platform. The simulated edges have a length of approximately 41 nm, a thickness of around 4 nm, and the angle at the vertices of ∼ π/4, which agree well with our design parameters.

### AFM Imaging

The planar origami structures were characterized by AFM in air. The samples prepared were deposited on mica surfaces for measurements. The target sample was diluted to ∼0.5 nM in 1× TAE buffer at 6 mM Mg^2+^. A 10 μL aliquot of the diluted sample was added onto a freshly peeled mica surface and deposited for 5 minutes. Then the liquid was blown away by compressed air. Approximately 80 μL of DI water was added to the mica surface afterward and immediately blown away to avoid salt accumulation. AFM imaging was carried out with a Bruker Dimension Icon AFM using ScanAsyst-Air probes in the Peak-Force tapping mode. We followed the same procedure for sample preparation and measurement as previously described.^17^ The lengths of edges and the angles at vertices were measured and matched with our designs.

## RESULTS AND DISCUSSION

### Design of Architected Metastructures from DNA

**Figure 1**a shows the proposed auxetic design inspired by an Islamic geometric pattern. The Islamic pattern consists of periodic 2D arrays of large and small stars shown in blue and orange. The unit cell includes one small and two large stars which can slide along each other’s edges as depicted in Figure 1b. Upon loadings on horizontal directions, the sliding mechanism results in auxetic reconfigurations globally.^38^ This Islamic pattern design allows only one reconfiguration due to the limitations on sliding directions. By modifying the structure, we propose a new design which enables multiple reconfigurations with a single structure. To model the system, we simplified the geometric pattern to a design of sliding 4-point stars with identical sizes as illustrated in Figure 1c and 1d. The angles at four vertices of each star are designed to be π/4. Sliding of neighboring blue stars along the edges of the orange stars simultaneously in horizontal directions will result in a contraction of the structure both horizontally and vertically (Figure 1c). Note that the shrinkage in *x* and *y* directions will not be equal. By calculating the displacements in both directions, we can find the Poisson’s ratio:

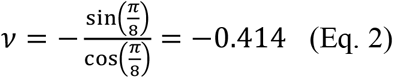

**Figure 1.**
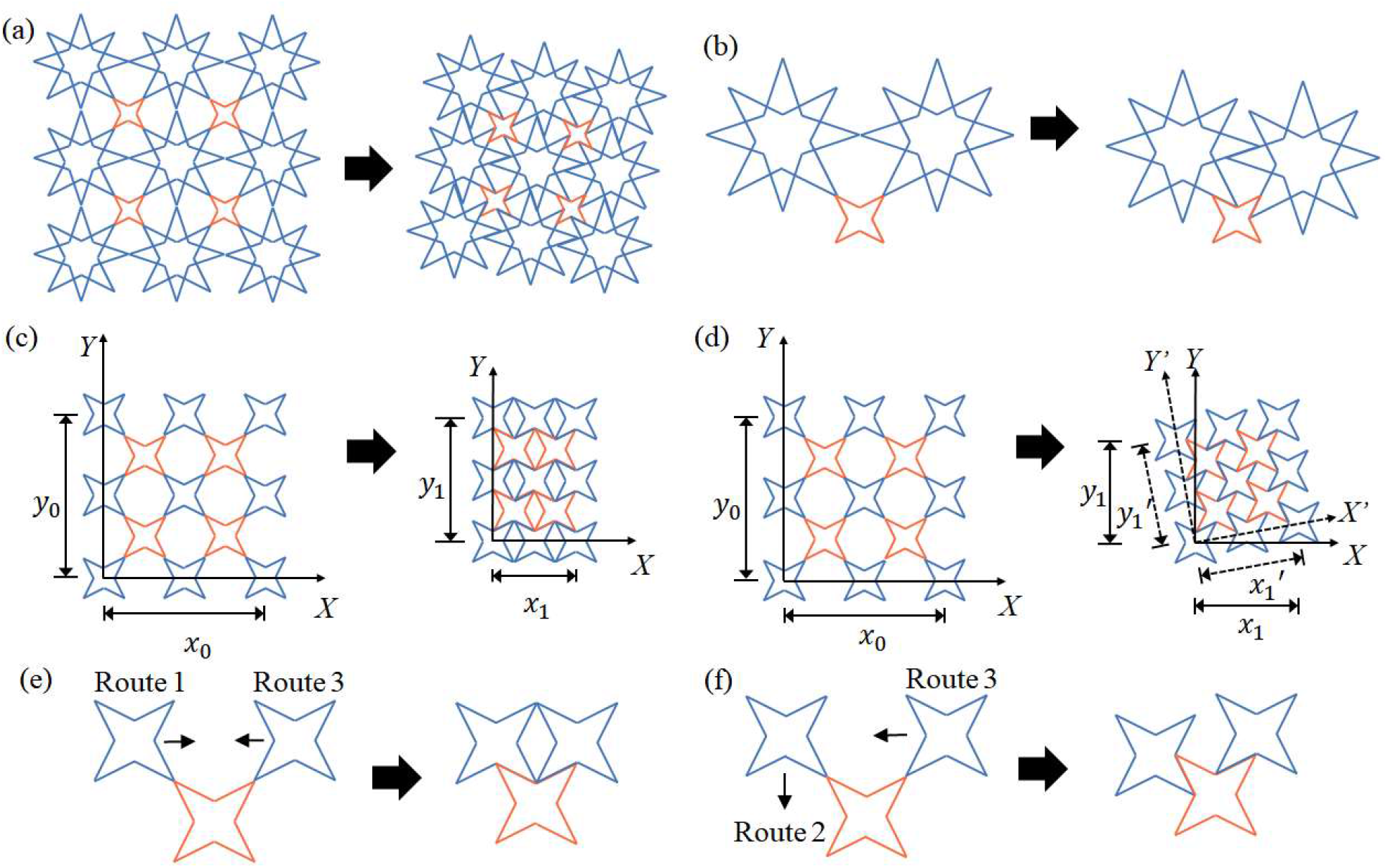
Schematics of the Islamic pattern and the auxetic design stuided in this work. (a) Periodic 2D cellular structure of the Islamic pattern. The large (blue) stars slide along the edges of both large and small (orange) stars, resulting in a global auxetic behavior. (b) Unit cell of the Islamic pattern with two large and one small stars. (c)-(d) A simplified design that consists of 4-point stars with identical edge lengths of 40 nm and angles of π/4. Horizontal sliding in unison between neighboring stars will lead to a contraction in both horizontal and vertical directions. This auxetic reconfiguration leads to a negative Poisson’s ratio of *ν* = -0.414. (d) If the adjacent stars move in different directions (e.g., one vertically and the other horizontally), the structure will demonstrate auxetic properties with *ν* = -1 via global rotation. By programming the sliding directions, two distinct auxetic behaviors can be demonstrated and structural mechanics may be tuned accordingly. (e)-(f) Unit cell that consists of two blue and one orange stars. (e) The horizontal movements of the left and right blue stars are named Route 1 and 3, respectively, resembling the structural transformation depicted in (c). (f) A vertical sliding of the left blue star is termed Route 2. Simultaneous Route 2 and 3 motions represent a centrosymmetric rotation illustrated in (d). Note vertical sliding in both stars will be redundant due to symmetry.

In contrast, a completely different structure will emerge if one of the two adjacent blue stars slides vertically, while the other moves horizontally. In this case, the entire structure will rotate for an angle of π/8, as shown in Figure 1d. This centrosymmetric movement will result in an auxetic behavior as well as a global rotation. Calculating the displacement in both *x* and *y* directions yields:

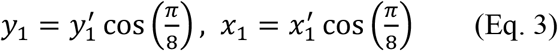

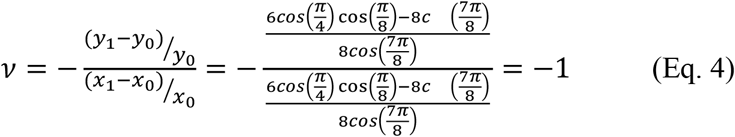

The rotation with *ν* = -1 is consistent with that of other centrosymmetric designs such as rotating squares.^17^ Overall, the auxetic properties of the sliding stars can be modulated by directing the sliding behaviors into a desired combination.

We further simplified the design into a unit cell of three (2 blue and 1 orange) stars which can be constructed by DNA origami (Figure 1e and 1f). Each edge of the stars is designed to have a length of approximately 40 nm long and the vertices have angles of π/4. These design parameters were later confirmed by measurements from oxDNA simulations and AFM imaging. Three sliding routes are made available for each unit cell. The horizontal sliding of the left blue star is named *Route 1*, while its vertical movement is termed *Route 2. Route 3* represents the horizontal sliding of the right blue star. Note that the vertical sliding of the right blue star is omitted due to symmetry. By programming the sliding behaviors, we can generate two combinations (Route 1, 3 or Route 2, 3) between the routes, which will give rise to two distinct auxetic properties. In a periodic cellular structure, the distinct Poisson’s ratios indicate the difference in mechanical properties.

In this work, a unit cell of the structure capable of the sliding behaviors is demonstrated with a wireframe DNA origami method. This strategy uses edges and multi-arm joints to represent geometric patterns.^39,40^ In wireframe origami, edges are composed of dsDNA bundles for structural integrity and are connceted by ssDNA at joints for flexibility. This method is efficient for material usage and allows for designing larger structures with a limited number of nucleotides (nt).^41^ To enhance the stiffness of the edges of the stars, we followed the principles previously described by Li *et al*.^17^ They demonstrated that edge thickness *t* must be sufficiently large for a given length *L* and ss-joints at vertices must experience a certain level of tension (termed joint stretch *η*) in order to avoid significant flexure or distortion during reconfiguration. Our edges and joints are designed with four duplex bundles (*t/L* ≈ 0.1) and a stretch level of *η* ≈ 55% to meet the design requirements. In addition, the edges are designed in a honeycomb arrangement to avoid internal strains in the structure as shown in **Figure 2**a. Here, the long blue arrows represent the routing of scaffold sequences, and the short gray lines denote staple strands. A three-dimensional molecular model from oxDNA is also shown in **Figure S1**. Given the design parameters of our proposed structure, single origami may not provide enough nucleotides. Therefore, we adopted a double scaffold strategy developed by Dietz and coworkers^42^ who used multiple orthogonal scaffolds with minimal interferences for larger origami structures. Figure 2(c)-(e) illustrates the routing of our origami design, where two kinds of scaffolds are differentiated by colors: 9072-nt scaffold (named 9k scaffold for simplicity) provided by the Dietz group is shown in blue color and commercially purchased 8064-nt scaffold (8k scaffold) is shown in orange color. The three-star structure is composed in a manner that each scaffold winds into one and a half stars, and the two scaffolds are connected by connecting staples to form the complete three stars. It is worth noting that the left one and a half stars in blue have a one-way connection given the linearity of the 9k scaffold, while a two-way connection is used for the right one and a half stars in orange as the 8k scaffold is circular. To differentiate between the left and right stars, a ss-loop of scaffold is designed on the top of the right star.

**Figure 2.**
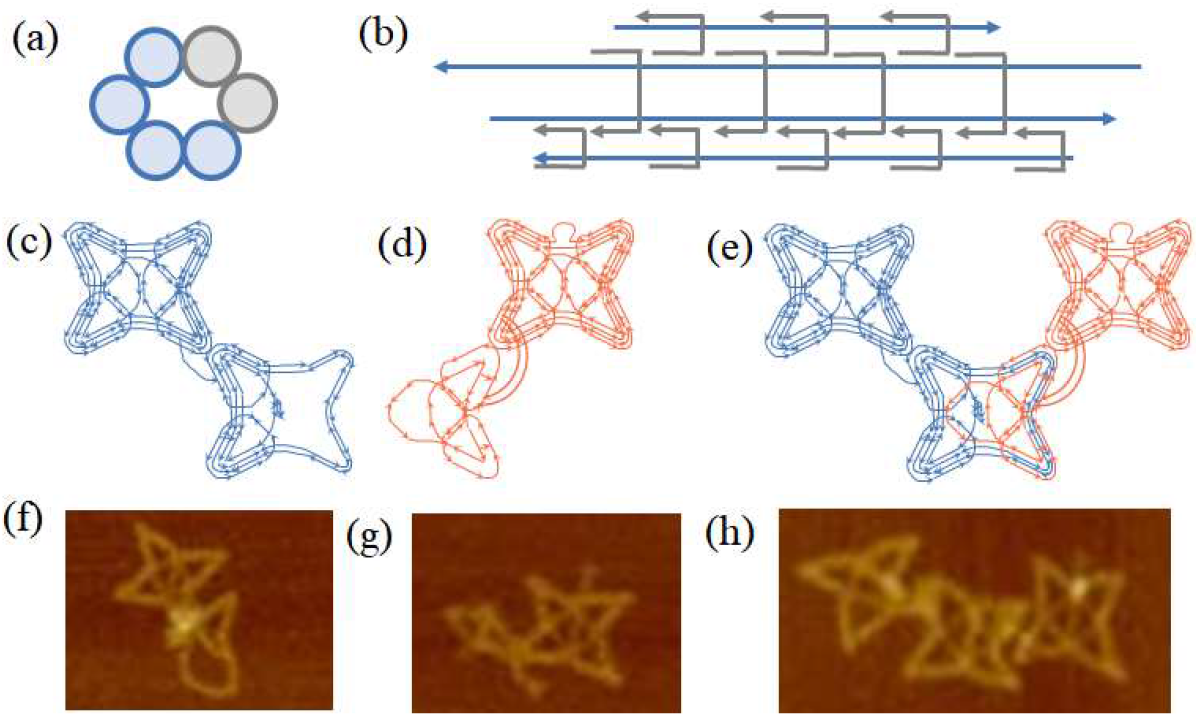
Unit cell design of three 4-point stars and experiment results with wirefram DNA origami. (a) Schematic of side view of arrangements of dsDNA bundles of an edge. Each edge consists of four bundles, and they are arranged in a honeycomb, where their locations are indicated with the blue color. (b) Schematic of front view of an edge segment. Each edge has four dsDNA bundles represented by four blue lines. They are connected by grey staple strands that go across different bundles to hold structural integrity. Routing of (c) 9k and (d) 8k scaffold strands. A tail of free 8k-scaffold loop is used for distinguishing between left and right halves. (e) Routing of the fully connected three stars. (f)–(h) AFM images of wireframe DNA origami. (f) Assembled half structure from the 9k scaffold and staple strands. (g) Origami half structure with the 8k scaffold and staple strands. (h) Fully assembled three stars using both 9k and 8k scaffolds. A tail of free 8k-scaffold loop is observed on the right star in (g) and (h).

AFM imaging characterized the assembled DNA units. The structures from a single scaffold (e.g., 9k or 8k strands) are tested initially with a scaffold concentration of 5 nM in 1× TAE buffer with 6 mM Mg^2+^. Both scaffolds formed half structures as designed; the 9k scaffold shapes the left star and half of the middle star (Figure 2(f)), while the 8k scaffold forms the right star and the other half of the middle star with a small loop at the top of the right star (Figure 2(g)). Finally, a whole structure assembled from both scaffolds is shown in Figure 2(f). Note that the right and left stars are connected with the middle star by unpaired segments of the scaffolds as indicated by 3 lines (one in blue and two in orange) in Figure 2(e). However, the relative positions between the stars are not determined (i.e., undefined position). Their locations are close to each other but random due to the deposition on mica surfaces – positional linker strands will determine the exact positions between the stars (*vide infra*). The structures are also examined with agarose gel electrophoresis. As shown in **Figure S2**, the half structures with the 8k scaffold or the 9k scaffold as well as the full three-star structures all have clear bands in the gel where the 8k-structure moves the fastest and the complete three stars were the slowest corresponding to their molecular weights.

### Demonstration of Auxetic Sliding Behaviors

With well-built structures, we designed sliding behaviors of the auxetic stars via toehold-mediated strand displacement as illustrated in **Figure 3**. For each route, the three stars are arranged to start with undefined position, where no binding is provided to determine relative positions between stars (state (i)). On one of two edges sliding against each other, three locations are modified with a 10-nt extension as binding entities (as shown in edge 1). Initially, the binding strands on the opposite edge (edge 2) are not provided, thus the stars remain in an undefined position as shown as state (i) in Figure 3(b). Then, each route can be programmed to take three positions to guide the sliding behaviors. To initiate the sliding, staples on edge 2 with toeholds are replaced by new staples (P1 staples in green color) via strand displacement. This allows the stars to initially bind to each other at the opposite vertices (position 1, state (ii)). Next, the binding sequences (shown in green) can be replaced with new sets of linker strands via toehold mediated strand displacement to direct the edges to overlap halfway where the stars slide more into each other (position 2, state (iii)). The binding staples for position 2 are shown in red color. Finally, with the same strand displacement mechanism, the edges can be programmed to slide to fully overlap with each other resulting in a configuration of position 3 (state (iv)). The staple designs for position 3 are shown in purple color. With the strand displacement mechanism used, this process may also be reversed to direct the stars to move back from position 3 to position 2 and then to position 1.

**Figure 3.**
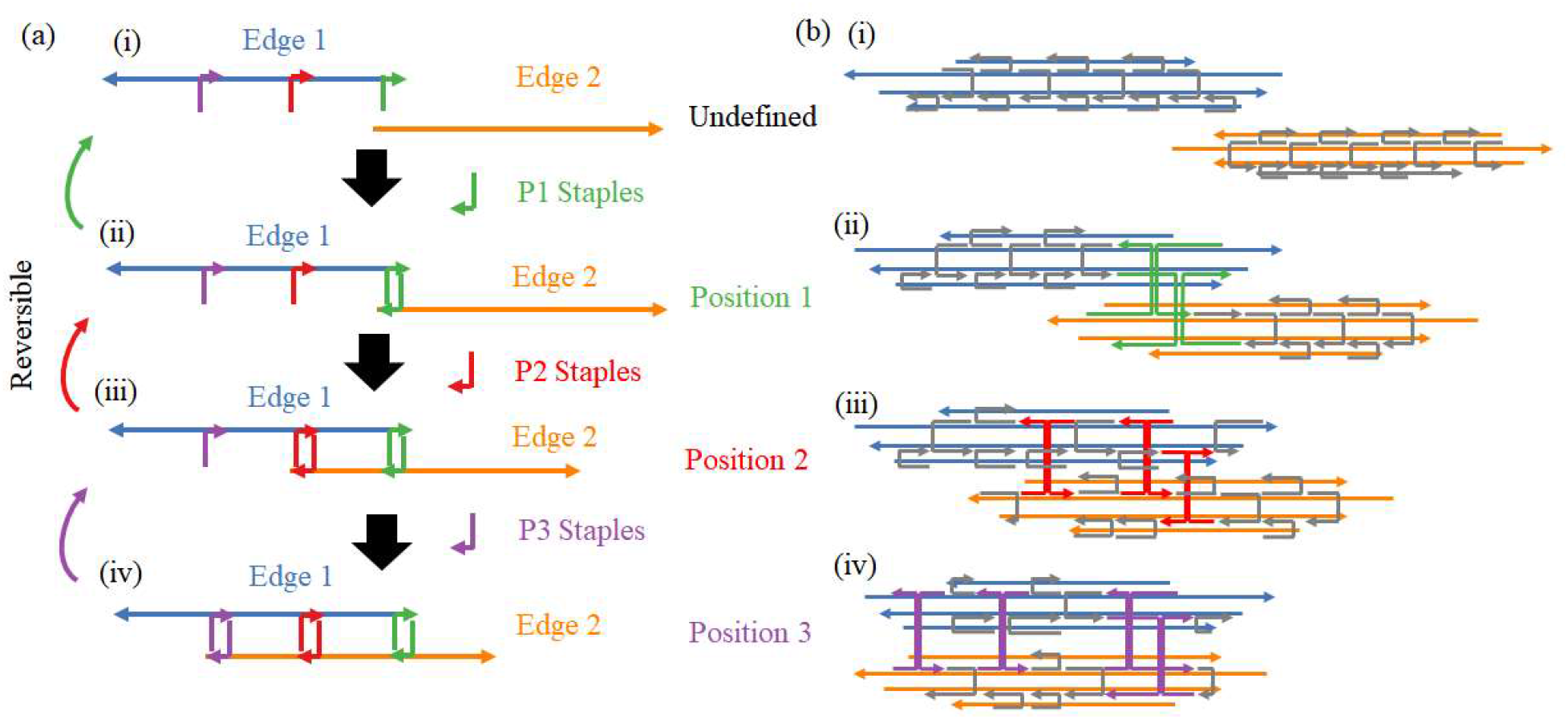
Illustration of sliding mechanism. (a) Two opposite edges (edges 1 and 2) on two stars sliding against each other are shown in blue and orange colors starting from an undefined conformation (i.e., edges are not associated). For sliding, staples on edge 2 are replaced via a strand displacement mechanism (see the text for details). Initially, a set of staples on edge 2 are replaced by P1 staples (shown in green), guiding the edges to position 1 (state (i) ⟶ state (ii)). By sequentially replacing the staples to P2 (red) and P3 (purple) staples, the edges can slide to position 2 and 3 (state (iii) and state (iv), respectively). This process may be reversed by replacing the positional staples in an opposite order. (b) Staple design for the edges at undefined (unassociated) position (state (i)). Four blue lines and four orange lines represent blue and orange edges in (a), respectively. The blue and orange scaffold routes are consistent with those in Figure 2(e). The gray staples are the structural staples for integrity of the edges. The binding staples shown in green color are used for position 1 (state (ii)). Staple design for the edges at position 2 (state (iii)). The binding staples in red color associate with both edges and bring the edges to overlap. Staple design for position 3 (state (iv)). Purple staples are binding staples to guide the edges to slide fully into each other.

The sliding mechanism is firstly tested on individual routes to ensure independent sliding beahviors. All routes start from the undefined positon as presented in Figure 2(e) and (h). On the edges, staples used for determining the relative postions are designed with a 10-nt toehold. The reconfigurations take place in two steps: (1) previous staples are displaced by adding complementary releaser strands. These strands bind to the staples defining positions on the edges and remove them. (2) The undefined structures are reannealed with new positional staples. These new staples relocate the edges to a new relative position as designed.

For each route and each position, we performed MD simulations using oxDNA to verify the formation of the structure. **Figure 4** presents the computed structures and corresponding, reconfigured origami units from AFM imaging. Figure 4(a)-(c) shows the sliding movement on route 1, where the left star starts from the tip of the vertices (position 1) and moves to the midpoint (position 2) and finally to the center of the middle star (position 3) as the staples are replaced sequentially. Route 2 sliding is demonstrated similarly as shown in Figure 4(d)-(f). The left star on route 2 slides vertically along the edge of the middle star. Starting from the corner (position 1), the star slides to a lower position (position 2) and reaches the destination at the junction in the center (position 3). Figure 4(g)-(i) shows the sliding on route 3, where the right star begins at the correct position (position 1) and then moves horizontally from right to left, reaching the middle point and the center point in the end (positions 2 and 3, respectively). More experimental results are included in **Figure S3**. For all the positions, both the experimental and simulation results show correct formation of the structures, and the relative positions of the stars are placed as designated.

**Figure 4.**
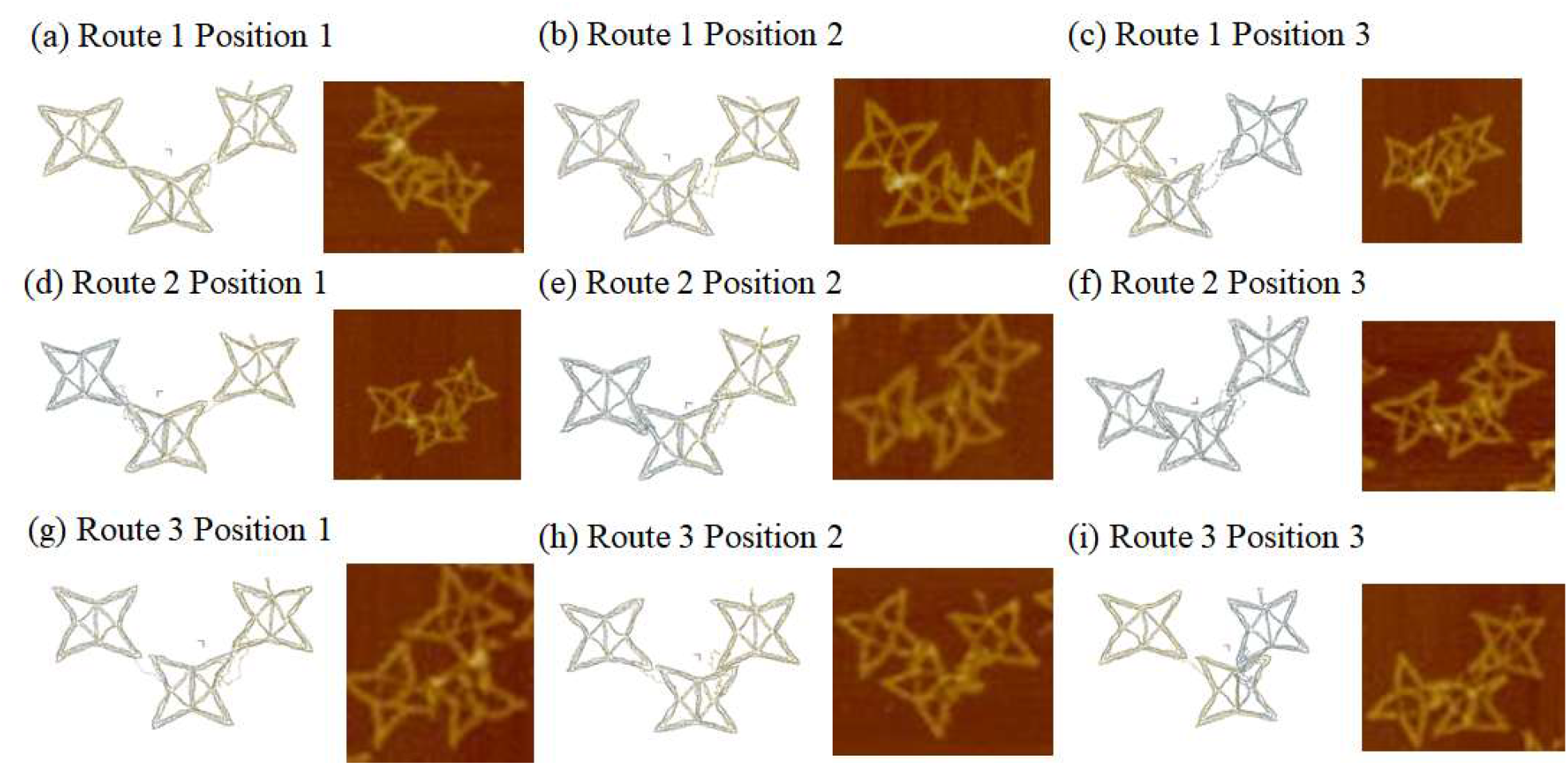
OxDNA simualtions and AFM images of nanostar structure formations and sliding behaviors of individual routes. For simulations and experiments, all structures start from undefined positions and the sliding behaviors are demonstrated at each position on individual routes. (a) Route 1 at position 1. (b) Route 1 at position 2. (c) Route 1 at position 3. (d) Route 2 at position 1. (e) Route 2 at position 2. (f) Route 2 at position 3. (g) Route 3 at position 1. (h) Route 3 at position 2. (i) Route 3 at position 3. The simulations and the AFM images confirm the correct sliding of the three-star structure programmed by strand displacement.

To generate tunable auxetic motions, we have explored the route combinations. As demonstrated above, the right star can slide in horizontal directions while the left star moves to two different routes upon the addition of replacement sequences (designed for intended routes). If routes 1 and 3 are selected, for example, the unit cell will translate into a Poisson’s ratio of -0.414. The combination between routes 2 and 3 will result in *ν* = -1. Here, the formation of the two combinations is confirmed both with oxDNA simulation and AFM imaging.**Figure 5**(a)-(c) present the combination between routes 1 and 3. Both left and right stars move horizontally along the edges to the middle star and meet in the center point (routes 1 and 3) with specific displacement sequences added. The unit cell shrinks in both horizontal and vertical directions. The reverse movement is also possible as shown in Figure 5(d)-(f). Two outer stars are directed to slide horizontally from the center (position 3) to sequentially move to position 2 and then to position 1.

**Figure 5.**
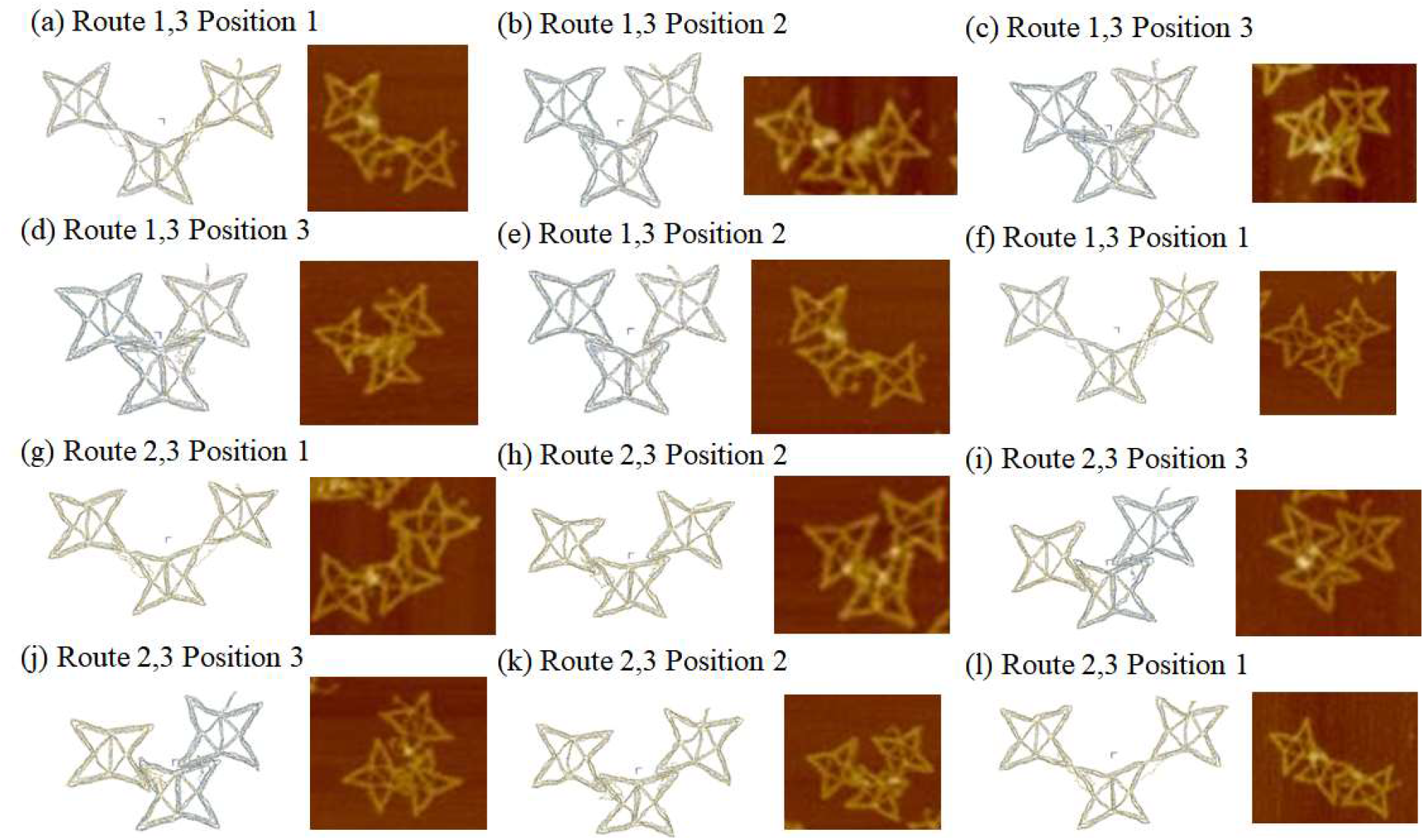
OxDNA simualtions and AFM images of combined auxetic reconfigurations by sliding stars. Sliding behaviors from vertices to center on route 1 and 3 are demonstrated at (a) position 1, (b) position 2, and (c) position 3. Outward movement from the center on route 1 and 3 are demonstrated at (d) position 3, (e) position 2, and (f) position 1. Sliding behaviors on opposite directions show reversiblity. Sequential movement from vertices to the center route 2 and 3 are shown at (g) position 1, (h) position 2, and (i) position 3. Sliding from the center to vertices on route 2 and 3 are demonstrated at (j) position 3, (k) position 2, and (l) position 1.

This results in expansion not only horizontally but also vertically, restoring the initial conformation. Figure 5(g)-(i) shows programmed sliding on route 2 and 3, where the stars start from position 1 and slide toward the center via position 2 and 3 sequentially. Similarly, Figure 5(j)-(l) shows sliding on route 2 and 3 from the center to the vertices going in a reverse direction from postion 3 to position 1. Additional experimental results for these combinations are shown in **Figure S4**. Since the sliding mechanism is based on the two step DNA reactions, it is possible to move directly to desired locations rather than sliding at sequential positions. Some examples of those positionings are demonstrated in **Figure S5**. The results demonstrate that our designed DNA origami can perform auxetic behaviors with tunable properties as well as reversibility as a one single unit cell structure.

## CONCLUSIONS

In this work, we proposed and demonstrated a novel design of metastructures with tunable auxetic properties by using DNA origami. In our design, a unit cell of three stars is arranged such that the left and right stars can slide along the edges of the middle star. Given the symmetry, three independent routes are used. The left star is designed to have two sliding orientations, horizontally and vertically, respectively as routes 1 and 2. The right star slides along the horizontal direction (route 3). A strand displacement mechanism is used to program the sliding behaviors. By using replacement strands, relative motions between the left and right stars can be directed in two combinations. Routes 1 and 3 will result in a Poisson’s ratio of *ν* = -0.414, while the combination of routes 2 and 3 will demonstrate a centrosymmetric rotation with *ν* = -1. In periodic structures, this can give rise to auxetic materials with tunable mechanical behaviors. Numerical simulations and experimental demonstrations show that our proposed origami structure can perform the designed sliding movement on each route individually. Combinations between the routes can be programmed with tunable auxetic behaviors with reversibility.

Given the biocompatibility and versatility, tunable DNA metastructures may be designed to interact with biological or chemical environments in a programmed manner; for example, as force-responsive sensors for biophysical studies and targeted drug delivery carriers.^26,30,43,44^ We envision that DNA-based structural design can be used to build metamaterials with multimode reconfigurability that can detect and respond to complex surroundings and show adaptive mechanical properties. For example, combined with aptamers, i-motifs or enzymatic reactions,^45-50^ DNA-based metamaterials will have strong potential to respond to environmental changes (e.g., pH change or the presence/absence of target molecules) with mechanical responses (e.g., changing stiffness, young’s modulus, etc). The strategy may be extended to various applications such as wound healing and vascular scaffolds. In addition, novel DNA materials may be developed as biological or chemical sensors, which respond to cues with reconfigurations or mechanical changes, thus opening new opportunities.

## ACKNOWLEDGEMENTS

The authors thank Dr. Hendrik Dietz, Max Honemann, and Michael Pinner at the Technical University of Munich in Germany for providing the 9072-nt scaffold strands. This work was supported by the U.S. Department of Energy (DOE), Office of Science, Basic Energy Sciences (BES) under award no. DE-SC0020673 (concept, design, and experiment) and the U.S. National Science Foundation under award no. 2025187 and 2134603 (computation).

## REFERENCES

1 Gibson, L. J. & Ashby, M. F. Cellular Solids: Structure and Properties. (Cambridge University Press, 1999).

2 Chan, N. & Evans, K. E. Indentation Resilience of Conventional and Auxetic Foams. Journal of Cellular Plastics 34, 231–260 (1998).

3 Scarpa, F. & Tomlin, P. J. On the Transverse Shear Modulus of Negative Poisson’s Ratio Honeycomb Structures. Fatigue & Fracture of Engineering Materials & Structures 23, 717–720 (2000).

4 Choi, J. B. & Lakes, R. S. Fracture Toughness of Re-Entrant Foam Materials with a Negative Poisson’s Ratio: Experiment and Analysis. International Journal of Fracture 80, 73–83 (1996).

5 Ali, M. N., Busfield, J. J. & Rehman, I. U. Auxetic oesophageal stents: structure and mechanical properties. Journal of Materials Science: Materials in Medicine 25, 527–553 (2014).

6 Jacobs, S. et al. Deployable auxetic shape memory alloy cellular antenna demonstrator: design, manufacturing and modal testing. Smart Materials and Structures 21, 075013 (2012).

7 Joseph, A., Mahesh, V. & Harursampath, D. On the application of additive manufacturing methods for auxetic structures: A review. Advances in Manufacturing 9, 342–368 (2021).

8 Budarapu, P., Yb, S. S. & Natarajan, R. Design concepts of an aircraft wing: composite and morphing airfoil with auxetic structures. Frontiers of Structural and Civil Engineering 10, 394–408 (2016).

9 Yanping, L. & Hong, H. A review on auxetic structures and polymeric materials. Scientific research and essays 5, 1052–1063 (2010).

10 Milstein, F. & Huang, K. Existence of a Negative Poisson Ratio in FCC Crystals. Physical Review B 19, 2030–2033 (1979).

11 Yeganeh-Haeri, A., Weidner, D. J. & Parise, J. B. Elasticity of α-Cristobalite: A Silicon Dioxide with a Negative Poisson’s Ratio. Science 257, 650–652 (1992).

12 Baughman, R. H., Shacklette, J. M., Zakhidov, A. A. & Stafström, S. Negative Poisson’s Ratios as a Common Feature of Cubic Metals. Nature 392, 362–365 (1998).

13 Schwerdtfeger, J., Heinl, P., Singer, R. F. & Körner, C. Auxetic Cellular Structures Through Selective Electron-Beam Melting. Physica Status Solidi (B) 247, 269–272 (2010).

14 Hengsbach, S. & Lantada, A. D. Direct Laser Writing of Auxetic Structures: Present Capabilities and Challenges. Smart Materials and Structures 23, 085033 (2014).

15 Subramani, P., Rana, S., Oliveira, D. V., Fangueiro, R. & Xavier, J. Development of Novel Auxetic Structures Based on Braided Composites. Materials & Design 61, 286–295 (2014).

16 Suzuki, Y. et al. Self-assembly of coherently dynamic, auxetic, two-dimensional protein crystals. Nature 533, 369–373 (2016).

17 Li, R., Chen, H. & Choi, J. H. Auxetic Two-Dimensional Nanostructures from DNA. Angewandte Chemie International Edition 60, 7165–7173 (2021).

18 Li, R., Chen, H. & Choi, J. H. Topological Assembly of a Deployable Hoberman Flight Ring From DNA. Small 17, 2007069 (2021).

19 Seeman, N. C. Nanomaterials based on DNA. Annual review of biochemistry 79, 65–87 (2010).

20 Rothemund, P. W. Folding DNA to create nanoscale shapes and patterns. Nature 440, 297–302 (2006).

21 Rothemund, P. W. K., Papadakis, N. & Winfree, E. Algorithmic self-assembly of DNA Sierpinski triangles. PLoS biology 2, e424 (2004).

22 Ke, Y., Ong, L. L., Shih, W. M. & Yin, P. Three-dimensional structures self-assembled from DNA bricks. science 338, 1177–1183 (2012).

23 Veneziano, R. et al. Designer nanoscale DNA assemblies programmed from the top down. Science 352, 1534–1534 (2016).

24 He, Y. et al. Hierarchical self-assembly of DNA into symmetric supramolecular polyhedra. Nature 452, 198–201 (2008).

25 Tikhomirov, G., Petersen, P. & Qian, L. Fractal assembly of micrometre-scale DNA origami arrays with arbitrary patterns. Nature 552, 67–71 (2017).

26 Marras, A. E., Zhou, L., Su, H.-J. & Castro, C. E. Programmable motion of DNA origami mechanisms. Proceedings of the National Academy of Sciences 112, 713–718 (2015).

27 Pumm, A.-K. et al. A DNA origami rotary ratchet motor. Nature 607, 492–498 (2022).

28 Du, Y., Pan, J. & Choi, J. H. A review on optical imaging of DNA nanostructures and dynamic processes. Methods and Applications in Fluorescence 7, 012002 (2019).

29 Du, Y., Pan, J., Qiu, H., Mao, C. & Choi, J. H. Mechanistic understanding of surface migration dynamics with DNA walkers. The Journal of Physical Chemistry B 125, 507–517 (2021).

30 Pan, J. et al. Mimicking chemotactic cell migration with DNA programmable synthetic vesicles. Nano letters 19, 9138–9144 (2019).

31 Sarraf, N., Rodriguez, K. R. & Qian, L. Modular reconfiguration of DNA origami assemblies using tile displacement. Science Robotics 8, eadf1511 (2023).

32 Song, J. et al. Reconfiguration of DNA molecular arrays driven by information relay. Science 357, eaan3377 (2017).

33 Dietz, H., Douglas, S. M. & Shih, W. M. Folding DNA into Twisted and Curved Nanoscale Shapes. Science 325, 725–730 (2009).

34 Ke, Y. et al. Multilayer DNA Origami Packed on a Square Lattice. Journal of the American Chemical Society 131, 15903–15908 (2009).

35 Snodin, B. E. K. et al. Introducing Improved Structural Properties and Salt Dependence into a Coarse-Grained Model of DNA. Journal of Chemical Physics 142, 234901 (2015).

36 Shi, Z., Castro, C. E. & Arya, G. Conformational Dynamics of Mechanically Compliant DNA Nanostructures from Coarse-Grained Molecular Dynamics Simulations. ACS Nano 11, 4617–4630 (2017).

37 Poppleton, E. et al. Design, Optimization and Analysis of Large DNA and RNA Nanostructures through Interactive Visualization, Editing and Molecular Simulation. Nucleic Acids Research 48, e72 (2020).

38 Lim, T.-C. A perfect 2D auxetic sliding mechanism based on an Islamic geometric pattern. Engineering Research Express 3, 015025 (2021).

39 Wang, X. et al. Planar 2D wireframe DNA origami. Science advances 8, eabn0039 (2022).

40 Zhang, F. et al. Complex wireframe DNA origami nanostructures with multi-arm junction vertices. Nature nanotechnology 10, 779–784 (2015).

41 Matthies, M., Agarwal, N. P. & Schmidt, T. L. Design and Synthesis of Triangulated DNA Origami Trusses. Nano Letters 16, 2108–2113 (2016).

42 Engelhardt, F. A. et al. Custom-Size, Functional, and Durable DNA Origami With Design-Specific Scaffolds. ACS Nano 13, 5015–5027 (2019).

43 Nickels, P. C. et al. Molecular force spectroscopy with a DNA origami–based nanoscopic force clamp. Science 354, 305–307 (2016).

44 Zhang, Q. et al. DNA origami as an in vivo drug delivery vehicle for cancer therapy. ACS nano 8, 6633–6643 (2014).

45 Cha, T.-G. et al. A Synthetic DNA Motor That Transports Nanoparticles Along Carbon Nanotubes. Nature Nanotechnology 9, 39–43 (2014).

46 Pan, J. et al. Visible/near-infrared subdiffraction imaging reveals the stochastic nature of DNA walkers. Science advances 3, e1601600 (2017).

47 Dong, Y., Yang, Z. & Liu, D. DNA nanotechnology based on i-motif structures. Accounts of chemical research 47, 1853–1860 (2014).

48 Zhang, Y. et al. Dynamic DNA structures. Small 15, 1900228 (2019).

49 Li, R., Madhavacharyula, A., Du, Y., Adepu, H. & Choi, J. H. Mechanics of Dynamic and Deformable DNA Nanostructures. Chemical Science 14, 8018–8046 (2023).

50 Liu, X., Lu, C.-H. & Willner, I. Switchable reconfiguration of nucleic acid nanostructures by stimuli-responsive DNA machines. Accounts of Chemical Research 47, 1673–1680 (2014).

