## Supplementary Information for "DNA Nanostar Structures with Tunable Auxetic Properties"

**Table of Contents**

S1. OxDNA Design of DNA Nanostars
S2. Agarose Gel Electrophoresis
S3. Additional AFM Images
S4. References
S5. DNA Sequences

**S1. OxDNA Design of DNA Nanostars**

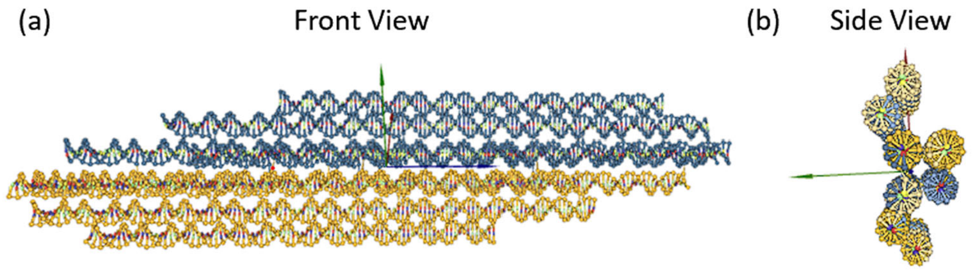


**Figure S1.** Schematic of edge design in oxDNA.[1,2] (a) Front and (b) side views of the edge design of four double-stranded DNA bundles. Dark blue and yellow colors indicate two distinct edges sliding against each other.

**S2. Agarose Gel Electrophoresis**

Agarose gel electrophoresis was used to examine correct formations of the DNA nanostructures. The agarose gel was prepared by dissolving 500 mg of powder in 50 mL of 0.5× TBE buffer (44.5 mM tris base, 44.5 mM boric acid, 1 mM EDTA and 11 mM magnesium chloride). The solution was heated to boil and then cooled down in a mold with addition of ethidium bromide (EtBr, 0.5 µg/mL) to form a rectangular shape gel with channels. Then, DNA samples were added into the channels and electrophoresis was performed for 2 hours under 60 V. The resulting gel was visualized with an UV transilluminator to identify bands of different samples. As shown in Figure S2a, individual structures of only 9072-nt or 8064-nt scaffolds have distinct bands. The half structures based on the 9k scaffold are slower than those with the 8k scaffold due to their larger molecular weight. With 4× staples per scaffold, the bands are stronger indicating better formations of structures. Therefore, 4× were used in the experiments. Figure S2b compares the effect of the ratio between the 9k and 8k scaffolds on the assembly of three-star structures. Different ratios between 9k and 8k scaffolds (1:1, 1.5:1, and 1:2 ratios of 9k:8k) demonstrate clear bands indicating the correct formation of three-star unit cells. Note that the 1:1 ratio shows a band of half structures from the 9k scaffold. In contrast, the other two cases do not show distinctly clear bands of half structures due to individual scaffolds. Given the gel results, we chose a 1:2 ratio between 9072-nt and 8064-nt scaffolds in sliding experiments.  It is noticeable that the assembled structures are faster than DNA ladders with corresponding molecular weights. This is attributed to the compact geometry of DNA origami enabling faster movement through porous agarose gel compared to free-form DNA.

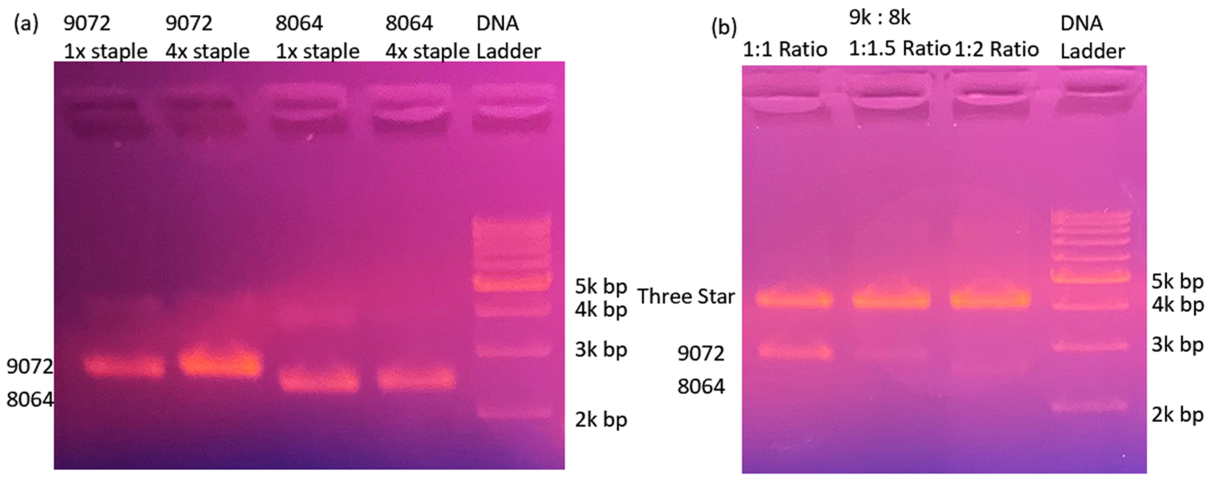


**Figure S2.** Agarose gel electrophoresis results. (a) Gel electrophoresis of half structures with 9072-nt or 8064-nt scaffolds with different staple concentrations. Two distinct bands of 9072-nt and 8064-nt based structures can be observed. The band with 4× staple concentration provides brighter bands. This condition is used for the formation of three-star structures. (b) Gel electrophoresis of three-star structures with various scaffold ratios between 9072-nt and 8064-nt scaffolds. It is obvious that the 1:2 and 1:1.5 ratios have stronger bands of three-star structures with weak signatures of individual half structures. Given the result, the 1:2 ratio has been used in experiments.

**S3. Additional AFM Images**

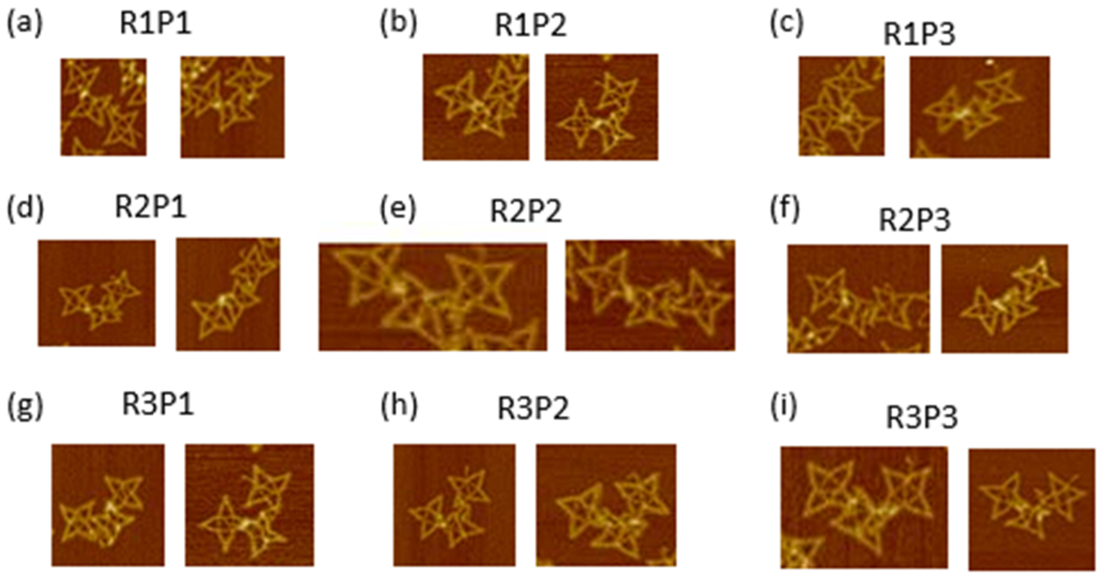


**Figure S3.** Additional experimental results for sliding on individual routes. The left star moves horizontally on route 1 at (a) position 1, (b) position 2, and (c) position 3. The left star moves vertically on route 2 at (d) position 1, (e) position 2, and (f) position 3. The right star moves horizontally on route 3 at (g) position 1, (h) position 2, and (i) position 3.

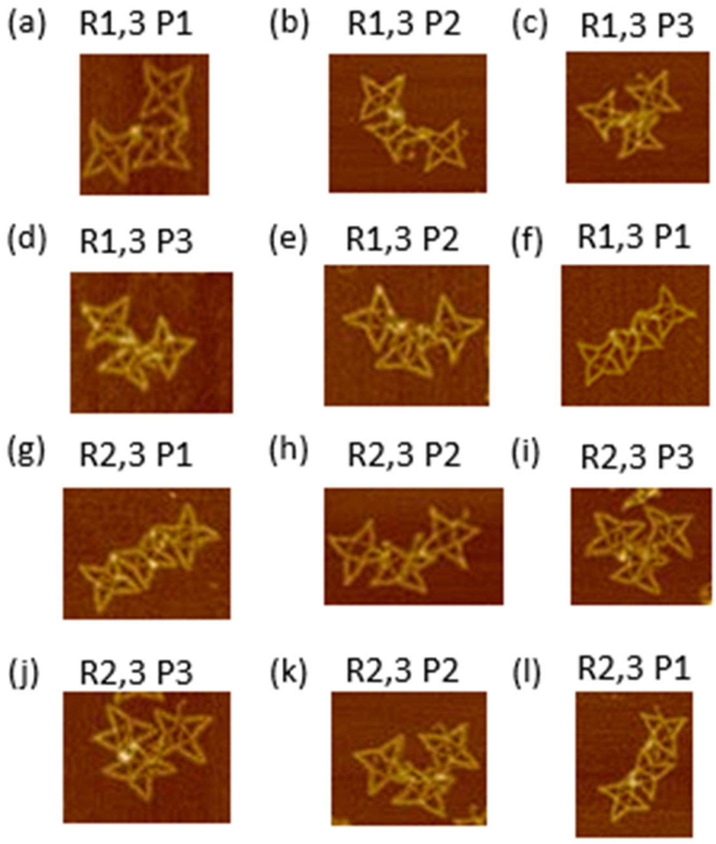


**Figure S4.** Additional experimental results for auxetic behaviors on combined routes. Auxetic sliding on combination between routes 1 and 3 are shown at (a) position 1, (b) position 2, and (c) position 3. Reversible movements are also demonstrated from (d) position 3 to (e) position 2, and then to (f) position 1. Similarly, for routes 2 and 3, reconfiguration is shown at (g) position 1, (h) position 2, and (i) position 3. Reversibly, sliding from (j) position 3 to (k) position 2 and finally to (l) position 1 is also demonstrated.

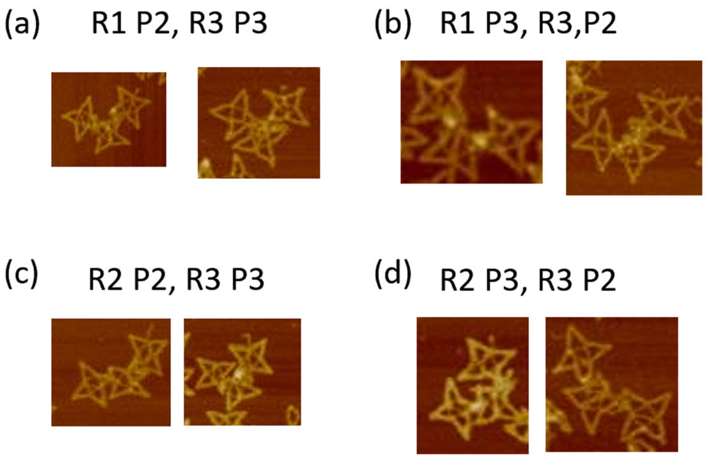


**Figure S5.** Additional experimental results for random positions. To show that control on individual routes is independent. Starting from undefined positions, several random reconfigurations are chosen. (a) Route 1 at position 2 and route 3 at position 3. (b) Route 1 at position 3 and route 3 at position 2. (c) Route 2 at position 2 and route 3 at position 3. (d) Route 2 at position 3 and route 3 at position 2.

### S4. References


1 Snodin, B. E. K. *et al.* Introducing Improved Structural Properties and Salt Dependence into a Coarse-Grained Model of DNA. *Journal of Chemical Physics* **142**, 234901 (2015).

2 Shi, Z., Castro, C. E. & Arya, G. Conformational Dynamics of Mechanically Compliant DNA Nanostructures from Coarse-Grained Molecular Dynamics Simulations. *ACS Nano* **11**, 4617-4630 (2017).

### S5. DNA Sequences

**Table S1.** Sequences of staple strands for the undefined position of three-star unit cells. Lowercase letters indicate toeholds and binding domains of positional staples.

| Name | Sequence |
|---|---|
| 0[139] | GTACTGTTGGTAACTTCCGCAGCACAACCACAAAAGTTCGGATGTCACC |
| 0[153] | AGAGATTTCGGCGTAGCTGGGGGAGTGTCTCTCGCAACAGAG |
| 0[69] | AAGTTCACCATGGTTAGGACCTAAGTGTTTATGCAACACGGA |
| 0[97] | CCCTACACGGCTAAACCGACTCTTGGCGTGTCTGTCTGAGAG |
| 1[129] | GGTGCAGCGAGGCGCAAGTAAGGATCGAGCTTGAGTGAGTGG |
| 1[154] | GTTGCCTTTTGGTTACCAGTAAGCACAC |
| 1[45] | TTTGGAGACTTCACTCATGACACGTTGACTATCCTTAAGAGT |
| 1[70] | CCCACCCCGCACTAGGGCTTACATACTGACACTACCTGGGAT |
| 1[84] | CATGGATCGGCTTGCTATGGTGACAAAAGTTTAAACGAATAT |
| 10[139] | CGCCGACTCCGCGATCACCGCCCATTAGGAGCCGTGGTAATT |
| 10[162] | ACGATATCCTCTATGTGAAGAATTCGGGACAACCGAC |
| 10[97] | GCTCTAGAGCGATCCGTCCTACCTGTCCCCACGAAAAAGATC |
| 100[139] | ACGAGTAGTAAATTCACCACCTTTTCGGTCATAGCTCACCGG |
| 100[153] | CGAGAAACAGAGCCTGTAGCGCGTTTTCACGTACAGCG |
| 100[90] | GTGAATTACAACGCCTGTATTTCATAATCAACTTTAATCATT |
| 101[126] | AAACAATCGGCGAAATCGGCAGGAACCGCCTCCCTCACCAGA |
| 101[150] | CCATGTTTACCGTCAGACGCCACCC |
| 101[171] | TTTGTGAGAGAAGAATCAAGTTTGCCTTTAGCAGTCCCGGAA |
| 101[91] | GGTGGAGCCGCCACGGGAACGGATAACCCCCCTTAACCGGAA |
| 102[104] | TTGCCATCTGCATTCCACAGACAGCCCTGCCAAGCTTTCAGA |
| 102[62] | CGATCTACGTTGTAAAACGACGGCCAGTCATAGTTCACCAGT |
| 103[119] | CCAGAGCGGGCTTGAGATGGTTTAATTTCAAAATCTTAGCGT |
| 103[161] | TCAGAACCGCCATTTTTAGAGCCACCACCCTCAGAG |
| 103[57] | CACTGATGGCTCATTTTTTTTTGTCAGGA |
| 103[77] | ACAAACTACCTTATGCGATTTTAAGAACGTTTCGTAGCGTAA |
| 104[111] | ATGAGGAAGGGTAGTGCTAAACAACTTTGGCGAAAGGGGGAT |
| 104[155] | CGTTGGGAAAACAACAGTCTTTCCAGACGTGCCAGGG |
| 104[76] | ATGCCACTACGAAGGCGGGATAGGAACAACTAAAGGGAAGGG |
| 104[97] | ATTAAACGGGTAAAGCGAAAGTTCAGCGGAGTGAGCTCTTCG |
| 105[112] | GTGCTGCGGGATTTCAACGGCTACAGAGGCTTTTAAAACGAA |
| 105[129] | AGTTGGGTAACTAGTAAATGAATTTTCTGTATAAGGCGATTA |
| 105[147] | TTTTCCCAGTCACGTTTTGTCTTATTACAGGTAGA |
| 105[28] | ATTCGCCTCACGTTGAAATTTTTTTCCAAAAAAAGGCTCCAAAAG |

| | |
|---|---|
| 105[45] | CGCAACTGTTGGAATTGCGAATAATAATTTTTATTCAGGCTG |
| 105[63] | CGATCGGTGCGGGCAATAGAACGTCACCCTCAGCAATACGTA |
| 105[84] | CTATTACGCCAGCTCAACAGTACAGCATCGGAACGAGTTTCC |
| 107[140] | CTAACGGGAAAAATCTACGTTAAGAGGACTAAAGACTTTTTC |
| 108[59] | CAGAACCGCCAATTCATCAGTTGAGATTTAGGCCGCCACCCT |
| 108[80] | CACCCTCAGAAAATACCACATTCAACTAATGCTTAGTACCGC |
| 109[101] | CGCCAAAAGGAGTATCACCGTACTCAGGAGGTAGATACATAA |
| 109[122] | ATTACGAGGCACCGGAATAGGT |
| 11[101] | AGTGCAAAGGTCTGGCGGATGTTCTAGATTAA |
| 11[126] | GCAGTCTCGATGTTAGA |
| 11[143] | CAGCACCGTTACGTCGAAAGCCACTTACGGGTAAACAAC |
| 11[166] | TAACACTAAAGCAATTGTCGG |
| 11[28] | CACCTTATGC |
| 11[38] | TCAATATTCTCATATAGACTA |
| 11[59] | CATAGTTCCCGGTTAAGCGTAGCGATGTCCAA |
| 11[84] | GACCAAATTAGTGTGGA |
| 110[139] | CAACCTAAAACGAACGCCCACTTGCTTTCGAGGTGGAAGATC |
| 110[90] | ACCCCCAAACCAAAATAGCGATAGTTGCAACACTCATCTTTG |
| 111[126] | GCACTCCAGCCAGCTATCAGCGCATAACCGATATATTCGGTC |
| 111[49] | TTGGTGTAGTAAAAACTATCATAACCCTCAAAGTACAACGGA |
| 111[91] | GAGGGGACGACGACAGTATCGGCCTCAGAATTTCTATGACAA |
| 112[104] | CTTGATACCGAGAGGCTTTTGCAAAAGAGCATCTGCCAGTTT |
| 112[150] | TTGTATCGGTTTTTCCGGCACCGCTTCTGGTGGAGCCTTTAA |
| 112[62] | GGGTAATAGATGGGCGCATCGTAACCGTAGTTTTGCAGACGA |
| 113[119] | CAACCATAGAGGCAAAAGAATACACTAAGCCGACATAAACAG |
| 113[154] | GCTGAGGCTTTTTGGAGTTAAAGGCCGCTTTT |
| 113[39] | TAAGAGCAACTGTTTAGACTGGATAGTTTTAGATTTAGTTTG |
| 113[77] | CGATAAAGCGATTATACCAAGCGCGAAACGTTTACCCAGAGG |
| 114[132] | GCATTAGACGGGAGAATTAACTGAACACAGACAGTTAGCTGA |
| 114[162] | ACAGAGAGAATAACATTTTGGGGGATACATTTCGCAATCTCCGT |
| 114[69] | TCAGAGAAGTAATGGATCTACAAAGGCTCGCCATCAAAAATA |
| 114[98] | CAAAGTCAGAGGGTACAAAAGGAGAGGGTAGCTATTTCTGGCC |
| 115[129] | ACCCGTCGGATATGGTCAATAACTGATATTCAGCGAGTAACA |
| 115[147] | GGGAACAAACGGCGACCATTACGCGAGCTGAAAAG |
| 115[161] | GATTGACCGTATTGGGATAGGTCACG |
| 115[45] | ACCAATAGGAAATCAGGTCATTGCCTGAGAGTTCATTTTTA |

| | |
|---|---|
| 115[70] | ATTCGCGTTTGAGATGTAGGTAAAGATTATTGAGCGCTAATA |
| 115[84] | TTCCTGTAGCCAGCATGCCGGGTGAGAAAGGCCGGCCTGAA |
| 116[104] | TAAATTATTTCATCAACATTAAATGTGAACCGTTCCAAATCA |
| 116[34] | CTGGAGCAAACTTTTTTTAGAATCGATGAACGGTAA |
| 117[119] | CCATCAATACTGTTTAGCTATATTTTCAAAAAACAGGGAAGC |
| 118[101] | TAGGATTAGCGTCATACAGGCAAGGCAAAGAAAGAGAAGGAT |
| 118[59] | CGTCGAGAGGGGCATCAATTCTACTAATAGTAGGATAAGTGC |
| 118[80] | CAGTACCAGGCGTAGCATTAACATCCAATAAAGGGTTTTGCT |
| 119[122] | TTAGCAAAATTGAGACTCCTCA |
| 12[139] | GTCCCAGTCAAGCAGAGTGGATATTTGT |
| 12[31] | GGATGGCGTTTTTTTCTGAAGTGATG |
| 12[55] | TAGGGAGGCACACACGCTTACGAC |
| 12[97] | TTCGCGGCATGGAAAGTGGATTTCGCTA |
| 120[132] | GCCCAATGGGAGAATCAGAAAAGCCCCAAAAACAGGTTTCTT |
| 120[153] | AACCCACGCAAGGACATATGTACCCCGGAATTGTAAACGTTA |
| 120[69] | GAAAAGTAGCCTCAAGCAAATCGTTAACTTTGGCGCGGTTGC |
| 121[133] | ATATTTATTGATAAGCCTTTATTTCAACAAGAATTGAGTTAA |
| 121[154] | ATATTTTTGTCAATTAAAAATTTTTAGAACCCTTTTTTATTTTAAATGCAATGC CTG |
| 121[70] | GGTATGATGTGTTCGAGCATAAAGCTAAGAAGCCCTTTTTAA |
| 122[104] | TGCTCGTGGCTGGTAATGGGTAAAGGAAGATTGTATAAGCAA |
| 122[174] | TCGTAAAACTAGCAGTTAAAATTCGCAT |
| 122[83] | ACACTGGGCCGGGTCACTGTTGCCCTGCCATAAACGTACCAA |
| 123[112] | TGTAATACTTTTGCAATAAGAGCAAGAAACAATGAATGACCC |
| 123[57] | CAATAAAAGCAGATATTTTTTTCAAAGTTAC |
| 123[98] | AAACATTAATAGCAATAGCTATCTTACCATCGGTTATCCCTT |
| 124[101] | ATTTTTTGTTTTCGAAATCCGCGACCTGCTCCATAAGAAACG |
| 124[143] | AATAAACAGCCGCGCAGACGGTCAATCATAAGCCAGTTACAA |
| 124[164] | AGCCTAATTTGGGAACCGAACTGACCAACTTTGTCTTTCCAG |
| 124[185] | CGCTAACGAGCGAAAGAGGACAGATGAACGGTATCTTACCAA |
| 124[80] | ATGAAAATAGCATCATCGCCTGATAAATTGTGAACGTCAAAA |
| 125[122] | GCCGGAACGAGATATTATTTATCCCAATCCAAATGTTACTTA |
| 125[196] | GTACAGACCAGTTTTATCCTGA |
| 126[195] | AGGGCGACACGGAATCCCCGAAAAGTGC |
| 127[112] | TTAGAGCCGTCAATTCAAAAGAAT |
| 13[119] | GGGGTCCGGGTCCAGCGAAGTCAATAGCGTCGAGCTATGAGTCTAGCAT |

| | |
|---|---|
| 13[77] | GCGTTGTGCCCACCGTCTCAAGACGGAAATTGGATGTGGCCTATGATTC |
| 14[59] | CGGAATCTGGTGCGTCAGCTGTCGGCTATGAAGATGGGCAGC |
| 14[79] | ATAATGCCATCGTTACTGCC |
| 14[87] | CATAAACTTTGAATTAGCTACGATCGTAGACC |
| 15[104] | CGGACAAGATCTGTCTTGCGTGTATCACTCTT |
| 15[122] | CTGATACGCGAGGTATGATGAA |
| 16[132] | GCAACTCTCAGTACTACAGA |
| 16[153] | AGAGCAGCCCGCGAAGGACTCGAGGCGAGCATCTTACTCTGA |
| 16[94] | ATGATCCAGTATTCCAACCGGTGTGCGCCTACAAGGCCCTTG |
| 17[112] | ATTAAACTAAATAAAAATTGG |
| 17[119] | GTCGTCATCTACT |
| 17[132] | GGGTGTTAAGGCTTAGAATGAC |
| 17[154] | GCGAATAGCTGCTTTTTGAGTTTAGATCGGGCCTTTTTCCGACTAGCTATCT AACT |
| 17[161] | GACCTGATCTGCC |
| 17[174] | TCAAGGAT |
| 17[55] | TTAAATCCTTTTCTACTCGTACGCTAGCCGGTAGCG |
| 17[77] | CTACAGATCCTCG |
| 17[90] | TGACATTGGTCTCTCGGCCCTA |
| 18[104] | GCCTGTGGGACGCAGCTCGG |
| 18[126] | GCAATACA |
| 18[168] | GCCAAAGC |
| 18[181] | AGGCCTTACCGAA |
| 18[84] | CTCGGATC |
| 19[119] | GTCGGCAACATAGCTGTGGACGATCCCAGTTTTTGGTACCTC |
| 19[140] | AGAAGATTTCCGGTCCGCGGT |
| 19[57] | CCGACCCAGTCTTGTTTTTTTTACAGTTA |
| 19[77] | CAGGTGTCCCGGTAGGGGAACGCTATATACTGGAGACCTCCCCTTGGTG |
| 2[104] | TCATTGTGGACCTTGGAAGACGAAGCAGATCTAGTCCTCGGG |
| 2[34] | GCTTGCCTTATTTTTTTTTTTGACATACCTGTTCTAGTCA |
| 20[111] | CTAGACGACCCCGTTAGTATACCAGGATGTAAGCTTCGCATA |
| 20[155] | GCCCCATGTGGCCCAATTAATCATCGTGGCCTGCTAT |
| 20[76] | TCTCAGTCATGTCGCTCCATAAGGCGATTCGGTGATAGGAAG |
| 20[97] | TGGTCATTGCTATCCAACTGCACAGGTGATTAGACAGATGAG |
| 21[112] | CATTAATTGGATGATTCCTCTGCAAATCTAAGTTGCGGTCGC |
| 21[129] | CTGCGGCCATTTCATGGGACTTTAGACCACCCACACTCGTCC |

| | |
|---|---|
| 21[147] | GGACTATTCAAATTACCAAAAACGCTGATGACCGT |
| 21[28] | ATAGTGAGAGCGTGCGTCTTTTTTATAAACAGGAATCGAAGAACCC |
| 21[45] | AGACCGAGTGAATGCAGGTAATCCAGCAAATACTCTGTGATA |
| 21[63] | AGGACGGGCGGCAGGTATATTACAGGGAGAGATATAAGCGCA |
| 21[84] | CAAACTAATAGGAGTAAATTGCATAATCTACAGTCACTATCT |
| 23[140] | TGTCCGAGGATTTGTGTTTGCATCACGTCGCTAAAGATGAAT |
| 24[101] | GTAGCCTCGCTATGGATTCCAGGCAGCGGTAGCCTCCAAGAG |
| 24[59] | TCTAAGAATCCGAATGTTCACTAACCGAACCCAAATATAGTC |
| 24[80] | GACATCGGACGCAGACAGGGGCTGGAAGGCCGTATGTTGGTG |
| 25[122] | CACATACAAACACAAGAATGCT |
| 26[139] | GGGGCAGAAGGAGGACTGCATAGCCCGCCCAATATCAGACGA |
| 26[90] | TCAGGTTAGGGAACCTGCCAGCACGGGCGGCAATAGATTGAG |
| 27[126] | GCGCCTGCGTGCAGAAGTTTGATATACTTCTAACGACACTGC |
| 27[49] | TATCTTCGGGTAAGCTGGTCTGACAGTCTGAGGAGTATGAGC |
| 27[91] | ACCATCGTTCAGTTCGTTCGTAAATCTCCATTTCTGATTGAA |
| 28[104] | AAAGAAATCTAGACCGGCAGGCTTTTAGAATTAGAGTGTCTC |
| 28[150] | CGTCGGAGTTGAACACAGGACGTTATGTGTCGGGCATTCGAC |
| 28[62] | AGATTCTGTTAATGGAAGCCGCGTTCTCAAATACCCGCGAAG |
| 29[119] | CACCGCCAAGCATGCGAGCCAACGACCGTGATCCTCCCCAAC |
| 29[154] | TTTGCGGGGTTTTTTTAGCACTCAGCCTCCGCA |
| 29[39] | CAACGCAGTGAGGGCCCATGGCGGCCTTTAATTGCCGGCCGA |
| 29[77] | ACACCGGTTGCGTTTTAACATTGTAGCACCGTTAGATAAGTG |
| 3[119] | TCCCCTGCTGTAGATGCTCGTGTCCGTAAGGTATA |
| 30[132] | GCTAGGACAGAGAATGTCGCGGTATAATCAACTTCTCTAGTA |
| 30[162] | TACGAGCCCTTAGGGATATCCATTCTCCGGGCGAGTAGTAAACC |
| 30[69] | GATACCCCCAGGCTATTTCCTAGAGATCGCGCGTCGGCCACA |
| 30[98] | AGGGGCCTACCAACTAACCTTGTCACCAGAAAGTTGCAACCAG |
| 31[129] | GCCGCCGGAGCTTATAACTAGTAGGGCAAGATCTCATCGCTC |
| 31[147] | TCCGTTCTGTACTTAACAAATACCGGCGTCATAAC |
| 31[161] | CCTGAAACTACTTGGCATGATTGTAA |
| 31[45] | ATTCGTCGGGAACTACTAGCCTTGGCCGGTTTACAGCAACTG |
| 31[70] | ATTTGCCGGTACGGCAGGAGGCCGGACCGTCCATTCGGATGA |
| 31[84] | CTGCAGACATAGGTGCCTGTAACTGCGCATATTGTCGCTGA |
| 32[104] | CGTTGCATGTAGAAGGCGACGGGTTATACATGAACGACACGG |
| 32[34] | AACGTAAAAAGTTTTTTTTGGTCCTTCGGAAAAATT |
| 33[119] | TGGTGACCAAGTATTGTGTGGGGCCAACAAATTCCCTGACCT |

| | |
|---|---|
| 34[101] | TCAAATACGAACGCCGTACGTGTTCCCAGTCAACCACTAAGG |
| 34[59] | GTCACAGGGTCCAGGAAAGGGTCAGCGGAACGTTGTTAAGAC |
| 34[80] | GGTCGCCCCTATATTAGAATAAGCATCGGTATAACGGATCTA |
| 35[122] | GCAGTAGTAGCGTACTGCGGTT |
| 36[132] | AAGCTGGCAACCGTTCCAGGTGCTAGTTCACACCCGCTAATA |
| 36[153] | GAATTGACTTTACAATCCCACCGGCTACGGGAGTACGCATTC |
| 36[69] | TAACGAGTCCCATATACAGGTCCATCTGGACATCTCGAATAA |
| 37[133] | GTGCCGAATCTGTGTTACTTAGTACACAAGCAACCGCCTTTT |
| 37[154] | CATCCTCATTGTTCGTATAGGGACCAGTGACGGTTTTGAATGAGCGCACTTT GCCA |
| 37[70] | CTTGCAGACTAGTGGGATCCGGAAGATTTTCGTGCTATACAA |
| 38[104] | AGATAAAAGCTCGTTGTTGCTCCTGACGTGTATACGATACAT |
| 38[174] | GCGCCATAGTAGGCAAAGTATCGAAAAG |
| 38[83] | GTAAAAGGGCCACGCTAGTGGCATCTATCGAGGTTATGGGAT |
| 39[112] | CAGGCGGGAGGATTCGTCCAAAGTCATTTTCGGTCGTGCCAT |
| 39[57] | CACATTTTTGTGCGTTTTTTTTGTGAATATA |
| 39[98] | TGCCATTCAATCACGTGCAATATATCTGACTCGGGCTCCTAG |
| 4[101] | CGAACACTGGGAAGGCCAATTATGATACTCTCGAACGACGCC |
| 4[59] | CCATCTTCAGGCCCCATCTCCTGGCGCCTTAGCACTCGACTA |
| 4[80] | TTTGGTGTTGGCTTAGACTTACAGGATGTAACCTAGAGCCTC |
| 40[101] | GATGCAATCTGTCTTCGGCTGAAAGGTAACACTTCCTACGCA |
| 40[122] | CAACGAAACTGATGCGCGGAAACCTTCGTAGTTTCCTTGCCG |
| 40[143] | TCCGCAAGCGGCCCCGGGTGCTTGGCTGCTCCTATGGTGTGA |
| 40[164] | GGGGATCCAAAACCCATGCGTTCGGCAGTCCAAGACTTGCTC |
| 40[185] | CTAACCCAAGAACTCTCATGTACAATTCCAATACGTATGCCT |
| 40[80] | CATCGATTGATGAGCGAGATCCACCGACCTTCACCATGTCCG |
| 41[196] | GGAGAGCGCTAGCGTGCTTCTT |
| 42[139] | CGTTAAGTAGACGCGATAACCGTGATCGGATAGAATTCTTTC |
| 42[153] | TTCACCCCTTCTTCGTTAATAGTTTGCGCCGCGTGACCCACG |
| 42[69] | TCTTACCCAATACGGGTTCCCAACGATCTCATCCATAGTTGC |
| 42[90] | CGGGGCGCACATAGCAGAACTTTAAAAGTGCTCCTACGCTAA |
| 43[105] | AGGGCTTACCATCTGGCCCCAGTGCTGCAATGATACAACGTT |
| 43[154] | CTCACCGTTCGCCAAAGATTGTTGATGG |
| 43[45] | GTCTATTTCGTAAGGCGAGTTACATGATCCCCTCAGCGATCT |
| 43[70] | CTGACTCCAGCTCCGGATAATACCGCGCAAAACTCTCAAGGA |
| 43[84] | TGTAGATAACTACGGTCGTTTGGTATGGCTTCATTCCCGTCG |

| | |
|---|---|
| 44[132] | GTTGCCATTGCTACAGGCATCGTGGTGTCACGCTCATACGGG |
| 44[34] | CATGTTGTGCATTTTTTTAAGCGGTTAGCTCCTTCGGTCC |
| 45[119] | GACTCGCTACGGCCCAGCATTGGATCATTGGAAAACGTTCTT |
| 46[59] | AATTTTCCCTTTGGAGTAGTCAGTTTACAAGGGCTATTAATT |
| 46[76] | AATCGTCAGTCTCAGATACGTGGTCAATGAAGTCTGTGCGGT |
| 47[105] | GGCGTATATCAATATATGTGAGTGAATAACCTTGCTTCTGTA |
| 47[122] | GAAAGTAGGAGACAGTACATAA |
| 48[132] | CGATGTATACTCAAGCAGTGTTATCACTAAAGTATATATGAG |
| 48[90] | TTACTTTCACCAGCGTTTCTGGGTGAGCCTAATCGACCTCAC |
| 49[133] | TAAACTTGTTGGCCCCAAGTCATTCTGAGTTGAGATCCAGTT |
| 49[150] | TTACCAATGCTTCCGATCGTTGTCAGAAGTAAGGTCTGACAG |
| 49[49] | GTCATGACAGCAATCGACGCTCACTAGGAAAAACAGGAAGGCAAAAT |
| 49[87] | TTTAAATTAAATAATTCTCTTACGCACCAGCCCCTAGATCCT |
| 5[122] | GTCAGAAGCGGGACGTACCGTA |
| 50[104] | CACTGCAAATGAAGTTTTAAATCAATCTCATGGTTCTGTGAC |
| 50[62] | TGAAGCTGATTATCAAAAAGGATCTTCAGCTGTTAACGAGGC |
| 51[119] | TGGTGAGACCCACTCGTGCACCCAACTGTGCTTTTATGGCAG |
| 51[147] | GAATAGTGTATGCGGCTTTTTAGTTGCTCTTGCCCGGCGT |
| 51[39] | AAGCTCCTGTCGTTGAACCGTACTTTTTTGGCTGCGATCATTACGAA |
| 51[77] | AATGTACGGTGTCATGCCATCCGTAAGAATCTTCAGCATCTT |
| 52[139] | ATTTTGCGGAACAATTCGCCTACCTGGTTTATATGAAGATCC |
| 52[162] | TTAAAAGTTTGAGTAACCAAGTTGGTTCGATCAGTGGCGGGGTC |
| 52[90] | TCCTGATTACAGTAAGGGACACCACTTATTTGCAAGCAGCAG |
| 53[126] | TTTGATCTTTTCTAGAAGTCGGATTGCTTTGAATACATTATC |
| 53[147] | TGACGCTCAGTGGACCTTATTACAAAATCGCGCAG |
| 53[166] | AACTCATGTTAAGGGATTTTG |
| 53[45] | ACAAACCACCGCGGTGTATGGGTCTGTAATTAGATCCGGCAA |
| 53[91] | ATTACGCTCATACAACAGTACCTTTTACAATTATCATCATAT |
| 54[104] | GCCTGGGGCAGAAAAAAAGGATCTCAAGGGGGCATGAAACAA |
| 54[34] | TCAGAATTCAGTTTTTTTTTCTAGCTTTT |
| 54[62] | GGTTAATCTGGTAGCGGTGGTTTTTTTGCGCCTATACGTCAG |
| 55[119] | TAACGGAAGAAACCACCAGAAGGAGCGGATCGGGAAGATATG |
| 55[77] | ATGAATATATCAGATGATGGCAATTCATAGGTTTAAAGTCGG |
| 56[103] | AGTTAATTACATCAAGAAAACAAAATTAATTACACTGAGCAA |
| 56[59] | TCATTTGAATTCGAATTATTCATTTCAATTACTTTAACAATT |
| 57[122] | ATTTAATGGTTACTTTTTCAAA |

| | |
|---|---|
| 57[78] | AAGAAGATGATGAAACAATCATCTTCTGACCTAATATATTTT |
| 58[111] | TTTAGTAAATCATAAGAACTGTATTCTGTTAGTTACAACTGG |
| 58[139] | TTTGGATTATACTTGAAGGGTTTTTGTAGTTGCCGCTACGGC |
| 58[153] | TATAATCTCAAAATTCGTAGGTCTCAAGAGTATTTGGTATCT |
| 58[90] | CAAATTCTTAAATAAGAATAAACACCGGTCATATGCGTTATA |
| 59[112] | GGCCTAATTGAGAGATTACTAGATAATGCTGAAAAAAGCCTG |
| 59[126] | TACACTAGAAGAACTATGCTATAGAACCTACCATACTGATTG |
| 59[154] | GCGCTCTCCTACTGTATTTGCACGTAAAACAGATTTTTAGAAATTGCGTAGAT TTTC |
| 59[171] | GTTACCTTCGGTCTTGCACAGGTTGACGCGACGCTGAAGCCA |
| 6[132] | GGGGCCGCGGTAGTATCGCTTCGAACGGCTCATTTGGCCATG |
| 6[90] | TACATCAAACAATTAAGAAACCCTGCCACTAGTATATCCGCG |
| 60[62] | GTAAAGGTAACAGGATTAGCAGAGCGAGGCTTTATTGATAAA |
| 60[83] | TTATGCCGTATGTAGGCGGTGCTACAGAGTTCTTGAAGTGGT |
| 61[57] | AATACCTCAACAGTATTTTTTTAATTGAG |
| 61[77] | TAAGGCGTTACCAGTATAAAGCCAACGCGACCGTGAGCCTGA |
| 62[111] | GCGGTCACAGCAGAAGATAAAACAGTAGAAGAACTCAAACTA |
| 62[155] | AATCGCCATTGCTGGTTTACATGCCCGCATCGCCACT |
| 62[76] | GCTGAGAGCCAGCAATAGCCCTTCCATATGCTGAATGCACGA |
| 62[97] | CACCGCCTGCAACATAAAAATCATAGCGAACAAACCCGACCG |
| 63[105] | TCGTCTTGAGTCCAACCCGGTAAGACACGACTTATTGAAACC |
| 63[147] | GGCAGCAGCCACTGGCGACGAAATATCCAGAACAA |
| 63[28] | TAGGTCGATATAACACTTTTTTTTTAGCTAGGCGGCTAG |
| 63[45] | GCTGGGCTGTGCCTACTCTTTTCAAGTGACCTTTCGCTCCAA |
| 63[63] | ACCCCCCGTTCAGCCCCCAAGTAAAACATCGCCATGTGCCAC |
| 63[84] | CTGCGCCTTATCCGGTTACCGACCGAACGAACCACGTATTAA |
| 64[132] | GCGCCGGGAGTCATCGACAAGTCAGAGGGGCACGCGTAACTA |
| 65[140] | TCGGCCTATTTTAGTAATAACATCACTTGCCTGAGAGGTGAG |
| 66[59] | ACAAAGAACGCTACCGCCAGCCATTGCAACAGCAATCGCAAG |
| 66[76] | GCAAATCGAAAAACGCTCATGGAAATACCTACATTTTGACGC |
| 67[105] | TCAATCGTAGGTTGGGTTATATAACTATATGTAAATGCTGAT |
| 67[122] | TCTGAAATGGAAACCTCCGGCT |
| 68[111] | AACCTCAGAACCCTGTCTCCCtggtgtgt |
| 68[139] | ATGAAAAATCTAAAGTAAGAATGAAAAC |
| 68[69] | CAAATCAACACGACCTGCTAAaaacctga |
| 69[112] | GTCCGCCGGCAACGTCTGACCTGAAAGCGCATCACCTTGCTG |

| | |
|---|---|
| 69[140] | GCGCTTTCTCAT |
| 69[152] | AGCTCACGC |
| 69[49] | GTTTCCCCTAGAGGCAGATTCACCAGTCACAGTTGAAAGGAA |
| 69[70] | CGTGCGCGGAGCGACAGTAATAAAGGGACATTCTGGCCAAC |
| 69[89] | ACCCTGCCGGCTGAGGTCGT |
| 7[133] | CCGTGGGTTGATCTTCTGAGTTACGGAACGTGTGACTAATCC |
| 7[150] | CTAAACGCGATTAGTGTCCTGACGGAGCACCAACATAATAAC |
| 7[49] | GCAGTGAGCCGTCCCCCCGTAGGGCATCGGTTAGGCTCGAAAGTCGC |
| 7[87] | CGGGGTCTCGTATGTCTCTTGATACCTATTTCTCTTAGGCAC |
| 70[104] | AACGCGCCTTACCGGATACCT |
| 70[125] | TAGGTTTTTTCTCCCTTCGGGAAGCGTGGAAAACATACGTGG |
| 70[146] | TTTGAAG |
| 70[160] | AATTATATCGCACG |
| 70[168] | GTGCGAGG |
| 70[62] | AATGTTTCCTGGAAGCTCCCT |
| 70[86] | GGTATCAATTTCTCCTGTTCCG |
| 71[140] | CACAGACAATATTTTTGAATGGCTTTTTGTCTTTAATGCGCGAACTG |
| 71[39] | TTTACATTGGTGTAAGGTTACACAACTTTTACCGGTCAAGAC |
| 71[98] | AGAGATAAATATCAAACCCTCAATCAATATCTGGTCAGTTGG |
| 72[125] | TTGACGGGGAAAGCCGGCGAAAGTGCCTATCCG |
| 72[162] | TAAATCGGAACCCTAATTTCACAGCGCG |
| 72[76] | AAAGGAGCGGGCGCTTAGGCCACATCGTAATGCATAAGGCCA |
| 73[105] | TCCGCCCCCCTGACGAG |
| 73[122] | cgtcctccgaccCATCACAAAAATCGACGCAATGTGTGAGAGCTCACAGTC |
| 73[154] | GCGAAACGCAACCCAGTACTCGAACATG |
| 73[161] | CCGACAGGACTTTAAAGATACCAGGC |
| 73[28] | CTTGCTAAGG |
| 73[38] | TGAGCAAAAGGCCAGCAA |
| 73[63] | GGAACCGTAAAAAGGCC |
| 73[80] | GCGTTGCTGGCGTTTTTCATTTCTGGTTGATT |
| 74[106] | CCTCAGTAACATAGGC |
| 74[118] | GTTTCGAGCTCACCTATCCTGCCGAGGGGATGCGA |
| 74[148] | AGACAGTGTTCAAGTCAGAGGTG |
| 74[34] | CGGAGACAAGGTTTTTTTCCTAAATGGGAG |
| 74[43] | TAAAGACGG |
| 74[55] | AGGGCTACTGAA |

| | |
|---|---|
| 74[83] | GCGGGGGAAGCGCCTGCACACGGGTGAGGAAAGCG |
| 75[140] | CCATCCAAGGGAGCCCCGATTTAGAGC |
| 75[98] | ACGTGCCCGTGGCGAGAAAGGAAGGGAATCCGTTGCGCCCAC |
| 76[101] | TAGATTAAGACCCTAACAAAAAGATACTGCAGAGCGATAGCT |
| 76[59] | ATCATAGGTCTCATCTGGGAGTTTCTCACGGAATTTATCAAA |
| 76[80] | GTCAATAGTGAAGGTACCCCTAGGCCGTTCCAGCTGAGAAGA |
| 77[122] | GACTTAAATGCCTTGAAAACAT |
| 78[115] | CCACACCCGCCGCGGCGTTAACTGAGCACAGTCGCGTAACCA |
| 78[136] | CGGTCACGCTGAATAAAATCTTAAAGGACTGAAATAACTGCAACTT |
| 78[153] | GGCGCTGGTCAAAACAAGTGG |
| 79[126] | CGAGCGGATGCACACGGGAGACCACGCCTCCACGAGCAAGTGTAG |
| 79[140] | TTGAGGTATCCTA |
| 79[153] | ACTCTGGATAACTG |
| 79[167] | GCCTCTGG |
| 79[56] | GCTCCAGATTTAT |
| 79[69] | CAGCAATAAACCAGCCAGCCGG |
| 79[91] | AAGGGCCCTCCATCAGAGCTAAATTCATGCGCTTAATGAACCATACTAA |
| 8[104] | TCGATAAGCACCTTTTGCTACTCAAGAATATTACGGAATAAT |
| 8[62] | TCTGCTGTTCCCGATGAAGCGTATAAAACGGGATTCCGGTTC |
| 80[104] | TATCCGCGAGCGCAGAAGTGGTCTGTCCTTAGAAC |
| 80[158] | CTAACGTGAGGCTTCTATA |
| 80[179] | GGCGTGAAGGCACGGCGCCCC |
| 80[74] | GCCGGGAAGCTAGAGTAAG |
| 81[161] | GTTCATTCATATTTTTTTGCCCTTCGAGGGCTGTTT |
| 81[57] | CAACATAGTAATTTATTTTTTTGTGGCAGCT |
| 81[70] | GCAGTTACCCTTACCTTTGTCTTACAAATGTAAATGTCTAGT |
| 81[84] | GGTGATCCAGTCTATTAATTGTT |
| 82[115] | AGACCGAGATATAATAGAGAACCATCACCCTACGTCTATCAGG |
| 82[136] | TCCCTTATAAAAGATAATCAGGGTT |
| 82[157] | AATAGGCCGAAAGAAGTACAATATTATTGAAGATGTATTTAGA |
| 82[178] | AAATCAGCTCACAATTCGATACTCATACTCTTAATAGGGGTTC |
| 82[195] | CGCGTTAATCCTTTGCCCGAACGTTAAAAGGGAATA |
| 82[94] | TTGTTCCAGTTATCTTTATTTTTGGGGTCGAGCAACGTCAAAG |
| 83[109] | GCGATGGCCCCATGAGCATTGTCTACTACGT |
| 83[126] | GGATACATATTTGACATTTATACATTTGAGGATTTATCGGCAAAA |
| 83[151] | AAAATAAACACCTTTTTTAGACTTTACAAATTTTTTAACC |

| | |
|---|---|
| 83[172] | CGCGCACATTATGTTGAACAACTCGTATTAAAATTTTTGTT |
| 83[196] | CACCTAAATTGTTGTTAAAATT |
| 83[70] | TGGACTCGTGCCGTAGGTTATCTAAAATTGGAACAAGAGTCC |
| 83[88] | GGCGAAAAACATCAAGTGGAGCACTAACAACGGGTTGAGTG |
| 84[139] | GGAAACGCAGAATCGGCCAACGCGCGGGGAGAGGCAAACCTGAGACGAT |
| 84[153] | CAGAAGGCAAAAGAGCCGGGTTACCTGCGGCGCTTTCGCACT |
| 84[576] | ATATAACAGTTCATAAATATTCATTGAATCCAGTTTCATTCC |
| 84[597] | GGTGTCTGGACCCTCAAATGCTTTAAACAGTAACTAAAGTAC |
| 84[69] | CAGATATCACATTAGGTGCTGCGGCCAGATGCCAACGGCAGC |
| 84[97] | GTATTACCGCGCCCACTGCCCCTGCGCGCCTGTGCGTGGTGC |
| 85[129] | GGGTCATTGCAAGCCAGCGGTGCCGGTGCCCCAGCATCAGCG |
| 85[154] | CAATCCGATCAGATACTGGCATGATTAA |
| 85[45] | CAACCGCAAGAAATGCGGCGGGCCGTTTTCACTGGTCAGCAG |
| 85[70] | ACCGTCGACTCTGTATTGCGTTGCGCTCAATAGCAAGCAAAT |
| 85[84] | CATCCCACGCAACCAGTGTCAGCTTTCCAGTCGGGGGTTTGC |
| 86[104] | CCAGCGCAGCTTACGGCTGGAGGTGTCCCTGCATCTCGTGCC |
| 86[34] | GGTCATACCGGTTTTTTTTTCTGCCAGCAC |
| 87[119] | AGCTGCATTAATATAATAACGGAATACCAAACCGA |
| 88[101] | TAATGCCCCCTATAAAGGTGGCAACATATAAATATAAACAGT |
| 88[59] | GAAAGTATTAATCCTTATTACGCAGTATGTTATCTGAAACAT |
| 88[80] | GAACCTATTATGCAAACGTAGAAAATACATACGCCTATTTCG |
| 89[122] | AGAAACGCAAAAACAGTGCCCG |
| 9[119] | AGCTTCTGGCCGCTCTGGCATATCACGAGTTGGCGCACTGAG |
| 9[147] | CATCCGGCACGATAATTTTTTAGCAGCAAGGTGAAGACCT |
| 9[39] | CAGATCTACGGGAGTGGTTCTGGACTTTTCACACAGCCTACAATTAT |
| 9[77] | GTCAATCCTCCCTCAGACTCTACGAGGGATTTCCAGATAGAG |
| 90[132] | TTCTAAGACAACATTTGAGgggctctg |
| 90[90] | GGAGGTTTTGAAGCCTTAAATCAAGATTACAATCAGACAAAA |
| 91[122] | AGAGCACATCCTCATAACGGAatggacca |
| 91[143] | ACGTGCCGGACTTGTAGATTCTTCGgcactagt |
| 91[49] | AGTTAAAATTCAACAAGTTTATTTTGTCAGTTGCTATTTTGCACCCA |
| 91[80] | ttgcgctcACGCGGTCCGTTTTTTCG |
| 92[104] | TTCGTAATCTCGTCGCTGGCAGCCTCCGGCCaaggtttc |
| 92[127] | GATCCCCGGGTACCGAGCTCACAaagtagac |
| 92[153] | CGTCCGTGAGCCTCCTCACAGACGAGCCGGAAGCAAGGCTTATCCGGTA |
| 92[169] | cgtgcgacGCGTGCCTG |

| | |
|---|---|
| 92[62] | GGGCGACCGATGCTGATTGCCGTTCCGGCAAtcgaactc |
| 92[85] | aacgctacGTTTACCAGCGCCAAAATAGAAA |
| 92[97] | TCATGGTCATAGgttggctg |
| 93[119] | ATTCCACAACGCGAGGCGTTTTAGCGAAGTTATCCGCTCGAA |
| 93[147] | TAAAGTGTAAAGCCTGTTTTTCCTAATGAGTGAGCTAACT |
| 93[39] | ACCACGGAATCGATTGAGGGAGGGAATTTAAATATTGACGGAAGGCG |
| 93[77] | ATTCATATGCTGTTTCCTGTGTGAAATTCCTCCCGACTTGCG |
| 94[139] | TAATCTTGACAAGAGATTGGCGTCACCGACTTGAGTGTGTAC |
| 94[162] | GGCTGGCTGACCTTCAACAAATACATTAAAGGTGAATATCCCGT |
| 94[90] | CTGCTCAGAGCCGCACCAGTAGCACCATAAACTTAAATTTCT |
| 95[126] | ATCGACATAAAAAATATCACCCTTGATATTCACAATCAAGAG |
| 95[147] | AAAAAAAGCCGCACAATTATTAATCCTCATTAAAG |
| 95[166] | GCCTTTTGTGATGAAGGGTAA |
| 95[45] | TCACGGAAAAACGTCACCAATGAAACCATCGAGGGATAGCTC |
| 95[91] | GCTCATTCAAAATCCGCCAGCATTGACATCAACGTAACAAAG |
| 96[104] | GAGCCAGTGCCGCCAGCAGTTGGGCGGTCCATTTGGAGGCAG |
| 96[34] | TAGCAGCACCGTTTTTTTCAGTAGCGAC |
| 96[62] | CCGGAAAGAGACGCAGAAACAGCGGATCTACCATTCAGAACC |
| 97[119] | GTCAGACACCGGATATTCATTACCCAAAGGAGGTTGGAATTA |
| 97[77] | ACCACCATTCAGTGAATAAGGCTTGCCCCCGCCACAGCAAGG |
| 98[59] | AGTTTTAACGGGAATGGAAAGCGCAGTCTCTGACTGGTAATA |
| 98[79] | ACAGGAGTGTAATTTACCGT |
| 98[87] | TTGATGATTCCAGTAAGCGTCATAGCAAGCCC |
| 99[104] | AATAGGAACTCATTTTCAGGGATACATGGCTT |
| 99[122] | CCCATGTACCGGAGCCACCACC |

**Table S2.** Sequences of replacement strands for sliding. The names are indicated in the order of route, position, and sequence number. Q represents complementary sequences

| Name | Sequence |
|---|---|
| R1Q.0.1 | GTCCAATAGCATAAGGTG |
| R1Q.0.2 | TAGTCTATATGAGAATATTGAATACTTCT |
| R1Q.0.3 | TTAATCTAGAACATCCGCCAGACCTTTGCACTTATATGAT |
| R1Q.0.4 | GCTCCACGTCCACACTAATTTGGTC |
| R1Q.0.5 | GATTCATCTCTAACATCGAGACTGC |
| R1Q.0.6 | GTTGTTTACCCGTAAGTGGCTTTCGACGTAACGGTGCTGTCGCAGGA |
| R1Q.0.7 | CATCACTTCAGAAAAAAACGCCATCCCAGCGCCG |
| R1Q.0.8 | CCCTGGTGGTCGTAAGCGTGTGTGCCTCCCTA |
| R1Q.0.9 | TAAGGTGTCCTTAGCAAG |
| R1Q.0.10 | TTGCTGGCCTTTTGCTCATCATCCTG |
| R1Q.0.11 | TTTGACGCGGCCTTTTTACGGTTCC |
| R1Q.0.12 | AATCAACCAGAAATGAAAAACGCCAGCAACGCGCAGTTTT |
| R1Q.0.13 | ACCCATACCTCGTCAGGGGGGCGGA |
| R1Q.0.14 | ACACATTGCGTCGATTTTTGTGATGGTTAACCG |
| R1Q.0.15 | CCGTCTTTATGCTGATG |
| R1Q.0.16 | ATGGCCATTTCAGTAGCCCT |
| R1Q.0.17 | TATGTTGTCGGATAGGCACTTTCGCCG |
| R1Q.0.18 | GCCTATGTTACTGAGGAGATCTGA |
| R1Q.0.19 | GAGAGGCTCGCGCTGTGAAATTAGGGTTCCGATTTA |
| R1Q.0.20 | CACCTCTGACTTGAACACTGTCTGGTGTGTC |
| R2Q.0.1 | AAGCTATCCGAGGATCTGTAG |
| R2Q.0.2 | TAGGGCCGAGAGACCAATGTCATCAACTGA |
| R2Q.0.3 | ACTGACTTAGTAGATGACGAC |
| R2Q.0.4 | GTCATTCTAAGCCTTAACACCCGTCGATGA |
| R2Q.0.5 | GGTACCAAGGCAGATCAGGTC |

| | |
|---|---|
| R2Q.0.6 | ATCCTTGATTTGATCG |
| R2Q.0.7 | TGGCTACCCCGAGCTGCGTCCCACAGGC |
| R2Q.0.8 | GATCCGAGAAGCGGGT |
| R2Q.0.9 | CCCTCTGATCTGTAGTACTGAGAGTTGC |
| R2Q.0.10 | TGTATTGCACTGGATC |
| R2Q.0.11 | GCTTTGGCATTCGTCA |
| R2Q.0.12 | TCGTACCCTTCGGTAAGGCCT |
| R2Q.0.13 | GCTATGGAATAAATCTGGAGC |
| R2Q.0.14 | CCGGCTGGCTGGTTTATTGCTGTTAATCGT |
| R2Q.0.15 | GCGGATAAAGTTGCAGTTATTGGACAGACAATGCGGC |
| R2Q.0.16 | ATAACGAACACTTCTGCGCTC |
| R2Q.0.17 | CAGCAATTCAGTTATCCAGAGT |
| R2Q.0.18 | CCAGAGGCACCGTCCG |
| R2Q.0.19 | CAGCCTGGAACAATTAATAGACTGGATCACCTTAGTAT |
| R2Q.0.20 | CTTACTCTAGCTTCCCGGCGCCTCCTA |
| R2Q.0.21 | GGGGCGCCGTGCCTTCACGCCCGATCATC |
| R2Q.0.22 | ACAAGTAACTCCCATTTAGGAAAAAAAACCTTGTCTCCG |
| R3Q.0.1 | ACGACCTCAGCCGGCAGGGTTCGGCGAT |
| R3Q.0.2 | GAACAACACGGAACAGGAGAAATTGATACC |
| R3Q.0.3 | ATTGTACTATGAGAAAGCGC |
| R3Q.0.4 | GCGTGAGCTCATAGAAC |
| R3Q.0.5 | TCAGGTTTTTAGCAGGTCGTGTTGATTTG |
| R3Q.0.6 | AGGGAGCTTCCAGGAAACATTATAGCTAT |
| R3Q.0.7 | ACACACCAGGGAGACAGGGTTCTGAGGTT |
| R3Q.0.8 | AGGTATCCGGTAAGGCGCGTTGTGAGACC |
| R3Q.0.9 | CTTCAAAAAGTCCCA |
| R3Q.0.10 | GGATTTATCGTGCGATATAATT |

| | |
|---|---|
| R3Q.0.11 | GCTTGCAACTAGCCGCCTAGCTAAAAAAAAAGTGTTATATCGACCTA |
| R3Q.0.12 | CCTCGCACACTTTCAT |
| R3Q.0.13 | CGAAAAAACGGACCGCGTGAGCGCAA |
| R3Q.0.14 | GAGTTCGATTGCCGGAACGGCAATCAGCATCGGTCGCCC |
| R3Q.0.15 | TGGTCCATTCCGTTATGAGGATGTGCTCT |
| R3Q.0.16 | ACTAGTGCCGAAGAATCTACAAGTCCGGCACGT |
| R3Q.0.17 | GAAACCTTGGCCGGAGGCTGCCAGCGACGAGATTACGAA |
| R3Q.0.18 | TTTCTATTTTGGCGCTGGTAAACGTAGCGTT |
| R3Q.0.19 | CAGCCAACCTATGACCATGA |
| R3Q.0.20 | CAGAGCCCCTCAAATGTTGTCTTAGAA |
| R3Q.0.21 | GTCTACTTTGTGAGCTCGGTACCCGGGGATC |
| R3Q.0.22 | TGGCTGACGTGCTGGCAGAAAAAAAAAACCGGTATGACC |
| R3Q.0.23 | CAGGCACGCGTCGCACG |
| R1.1.1 | CACCTTATGCccctccatagacaaactccg |
| R1.1.2 | gcctaccgagccgccgatcgTCAATATTCTCATATAGACTA |
| R1.1.4 | agggtcgtaagtAGTGCAAAGGTCTGGCGGATGTTCTAGATTAA |
| R1.1.5 | GACCAAATTAGTGTGGAcatcccatagcc |
| R1.1.6 | GCAGTCTCGATGTTAGAagctcgttcaaa |
| R1.1.7 | aggctggcttatCAGCACCGTTACGTCGAAAGCCACTTACGGGTAAACAAC |
| R1.1.8 | cgatcatcaacgagggggaaGGATGGCGTTTTTTCTGAAGTGATG |
| R1.1.9 | TAGGGAGGCACACACGCTTACGACtccacttctttcaggaaatc |
| R1.1.10 | CTTGCTAAGGgtctatggagggaaagcaca |
| R1.1.11 | tcatgcctcgatcggcggctTGAGCAAAAGGCCAGCAA |
| R1.1.12 | GGAACCGTAAAAAGGCCcccgtcgattgc |
| R1.1.13 | cgtcctccgaccGCGTTGCTGGCGTTTTTCATTTCTGGTTGATT |
| R1.1.14 | TCCGCCCCCCTGACGAGcccgtcgattgc |
| R1.1.15 | cgtcctccgaccCATCACAAAAATCGACGCAATGTGT |

| | |
|---|---|
| R1.1.16 | ggttgaagttccccctcgttTAAAGACGG |
| R1.1.17 | AGGGCTACTGAAgaaagaagtggaaagaaact |
| R1.1.18 | CGGCGAAAGTGCCTATCCGctccttcgaatg |
| R1.1.19 | ctaaaaagtcttCCTCAGTAACATAGGC |
| R1.1.20 | TAAATCGGAACCCTAATTTCACAGCGCGcgtcaagcctag |
| R1.1.21 | ataggcaacgctAGACAGTGTTCAAGTCAGAGGTG |
| R1Q.1.1 | cggagtttgtctatggagggGCATAAGGTG |
| R1Q.1.2 | TAGTCTATATGAGAATATTGAcgatcggcggctcggtaggc |
| R1Q.1.3 | ttaaaccgTTGGACATCGCTACGCTTAACCGGGAACTATG |
| R1Q.1.4 | CATCACTTCAGAAAAAAACGCCATCCttccccctcgttgatgatcg |
| R1Q.1.5 | gatttcctgaaagaagtggaGTCGTAAGCGTGTGTGCCTCCCTA |
| R1Q.1.6 | tgtgctttccctccatagacCCTTAGCAAG |
| R1Q.1.7 | TTGCTGGCCTTTTGCTCAagccgccgatcgaggcatga |
| R1Q.1.8 | CCGTCTTTAaacgagggggaacttcaacc |
| R1Q.1.9 | agtttctttccacttctttcTTCAGTAGCCCT |
| R1.2.1 | CACCTTATGC |
| R1.2.2 | TCAATATTCTCATATAGACTAgcaatcgacggg |
| R1.2.3 | ggtcggaggacgCATAGTTCCCGGTTAAGCGTAGCGATGTCCAA |
| R1.2.4 | ggcaaggcaagactttttagGGATGGCGTTTTTTTCTGAAGTGATG |
| R1.2.5 | TAGGGAGGCACACACGCTTACGACcattcgaaggagaggacggc |
| R1.2.6 | CTTGCTAAGGggctatgggatgtaaccaga |
| R1.2.7 | gaacagacacttacgaccctTGAGCAAAAGGCCAGCAA |
| R1.2.8 | TAAAGACGG |
| R1.2.9 | AGGGCTACTGAA |
| R1Q.2.1 | CATCACTTCAGAAAAAAACGCCATCCctaaaaagtcttgccttgcc |
| R1Q.2.2 | gccgtcctctccttcgaatgGTCGTAAGCGTGTGTGCCTCCCTA |
| R1Q.2.3 | tctggttacatcccatagccCCTTAGCAAG |

| | |
|---|---|
| R1Q.2.4 | TTGCTGGCCTTTTGCTCAagggtcgtaagtgtctgttc |
| R1Q.2.5 | gactgcacgcaatcgacgggGGCCTTTTTACGGTTCC |
| R1Q.2.6 | AATCAACCAGAAATGAAAAACGCCAGCAACGCggtcggaggacggtgtacta |
| R1Q.2.7 | cgtgctaccattcgaaggagCGGATAGGCACTTTCGCCG |
| R1Q.2.8 | GCCTATGTTACTGAGGaagactttttaggatagtcc |
| R1.3.1 | agcgttgcctatGGATGGCGTTTTTTCTGAAGTGATG |
| R1.3.2 | TAGGGAGGCACACACGCTTACGACctaggcttgacg |
| R1.3.3 | CTTGCTAAGGtttgaacgagct |
| R1.3.4 | ataagccagcctTGAGCAAAAGGCCAGCAA |
| R1.3.5 | GGAACCGTAAAAAGGCCggctatgggatg |
| R1.3.6 | acttacgaccctGCGTTGCTGGCGTTTTTCATTTCTGGTTGATT |
| R1.3.7 | CGGCGAAAGTGCCTATCCG |
| R1.3.8 | CCTCAGTAACATAGGC |
| R2.1.1 | CTACAGATCCTCGcgtcggtcgata |
| R2.1.2 | ttgttcgtaagaTGACATTGGTCTCTCGGCCCTA |
| R2.1.3 | GTCGTCATCTACTaggggaatcggc |
| R2.1.4 | actaccttggagGGGTGTTAAGGCTTAGAATGAC |
| R2.1.5 | GACCTGATCTGCCggcagactgttttcgcagca |
| R2.1.6 | ataacgcactatattttgatTCAAGGAT |
| R2.1.7 | GCCTGTGGGACGCAGCTCGGgtcacaccgggc |
| R2.1.8 | gagtccgccctgCTCGGATC |
| R2.1.9 | GCAACTCTCAGTACTACAGAtcctcgacgggg |
| R2.1.10 | gggagcactgcgGCAATACA |
| R2.1.11 | accaacatcagcctcaagctGCCAAAGC |
| R2.1.12 | AGGCCTTACCGAAgttgaatttacccataatca |
| R2.1.13 | GCTCCAGATTTATgcgtccccgtgg |
| R2.1.14 | attctctatgtcCAGCAATAAACCAGCCAGCCGG |

| | |
|---|---|
| R2.1.15 | ccaccttaaatgGTCTGTCCAATAACTGCAACTTTATCCGC |
| R2.1.16 | GAGCGCAGAAGTGcaccaattttcg |
| R2.1.17 | TTGAGGTATCCTAaaacgttt |
| R2.1.18 | ACTCTGGATAACTGaaacagtctgcccgattgcg |
| R2.1.19 | acggtgcgatcaaaatatagGCCTCTGG |
| R2.1.20 | ATACTAAGGTGATCCAGTCTATTAATTGTTatacaggggtaa |
| R2.1.21 | tattcgccaggcGCCGGGAAGCTAGAGTAAG |
| R2.1.22 | CTAACGTGAGGCTTCTATAgttggtat |
| R2.1.23 | cccgcgctagcttgaggctgGGCGTGAAGGCACGGCGCCCC |
| R2.1.24 | CGGAGACAAGGTTTTTTTCCTAAATGGGAGggtaaattcaac |
| R2Q.1.1 | tgctgcgaaaacagtctgccGGCAGATCAGGTC |
| R2Q.1.2 | ATCCTTGAatcaaaatatagtgcgttat |
| R2Q.1.3 | GCTTTGGCagcttgaggctgatgttggt |
| R2Q.1.4 | tgattatgggtaaattcaacTTCGGTAAGGCCT |
| R2Q.1.5 | aaacgtttTAGGATACCTCAA |
| R2Q.1.6 | cgcaatcgggcagactgtttCAGTTATCCAGAGT |
| R2Q.1.7 | CCAGAGGCctatattttgatcgcaccgt |
| R2Q.1.8 | ataccaacTATAGAAGCCTCACGTTAG |
| R2Q.1.9 | GGGGCGCCGTGCCTTCACGCCcagcctcaagctagcgcggg |
| R2.2.1 | GACCTGATCTGCCcgaaaattggtgggcgttca |
| R2.2.2 | ccggtaagcatttaaggtggTCAAGGAT |
| R2.2.3 | GCCAAAGCcagcatag |
| R2.2.4 | AGGCCTTACCGAActtaattt |
| R2.2.5 | TTGAGGTATCCTAgccgattccccttagccgca |
| R2.2.6 | ctttggccctccaaggtagtACTCTGGATAACTG |
| R2.2.7 | GCCTCTGG |
| R2.2.8 | atattaatcgcagtgctcccCTAACGTGAGGCTTCTATA |

| | |
|---|---|
| R2.2.9 | GGCGTGAAGGCACGGCGCCCCccccgtcgaggaaaaacgac |
| R2Q.2.1 | tgaacgcccaccaattttcgGGCAGATCAGGTC |
| R2Q.2.2 | ATCCTTGAccaccttaaatgcttaccgg |
| R2Q.2.3 | ctatgctgGCTTTGGC |
| R2Q.2.4 | aaattaagTTCGGTAAGGCCT |
| R2Q.2.5 | GCGGATAAAGTTGCAGTTATTGGACAGACcatttaaggtggaggttccc |
| R2Q.2.6 | aatcgccacgaaaattggtgCACTTCTGCGCTC |
| R2Q.2.7 | tgcggctaaggggaatcggcTAGGATACCTCAA |
| R2Q.2.8 | CAGTTATCCAGAGTactaccttggagggccaaag |
| R2Q.2.9 | TATAGAAGCCTCACGTTAGgggagcactgcgattaatat |
| R2Q.2.10 | gtcgtttttcctcgacggggGGGGCGCCGTGCCTTCACGCC |
| R2.3.1 | GACCTGATCTGCCccacggggacgc |
| R2.3.2 | gacatagagaatTCAAGGAT |
| R2.3.3 | gcctggcgaataGCCAAAGC |
| R2.3.4 | AGGCCTTACCGAAttacccctgtat |
| R2.3.5 | GTCTGTCCAATAACTGCAACTTTATCCGC |
| R2.3.6 | GAGCGCAGAAGTG |
| R2.3.7 | TTGAGGTATCCTAtatcgaccgacg |
| R2.3.8 | tcttacgaacaaACTCTGGATAACTG |
| R2.3.9 | cagggcggactcCTAACGTGAGGCTTCTATA |
| R2.3.10 | GGCGTGAAGGCACGGCGCCCCgcccggtgtgac |
| R3.1.1 | TAGGTCGATATAACACTTTTTTTTTAGCTAGGCGGCTAGcgtgtgcttgat |
| R3.1.2 | aatgtgaatctgGTGCGAGG |
| R3.1.3 | GGTCATACCGGTTTTTTTTTTCTGCCAGCACatcaagcacacgtgagagcc |
| R3.1.4 | gcagcgcacagattcacattGCGTGCCTG |
| R3Q.1.1 | ATCAAGCACACGCTAGCCGCCTAGCTAAAAAAAAAGTGTTATATCGACCTA |
| R3Q.1.2 | CCTCGCACCAGATTCACATT |

| | |
|---|---|
| R3Q.1.3 | GGCTCTCACGTGTGCTTGATGTGCTGGCAGAAAAAAAAAACCGGTATGACC |
| R3Q.1.4 | CAGGCACGCAATGTGAATCTGTGCGCTGC |
| R3.2.1 | GCGCTTTCTCATtcaatcggaagtatgtagac |
| R3.2.2 | aatagcagatacacccttgaAGCTCACGC |
| R3.2.3 | AACCTCAGAACCCTGTCTCCCcggttagtgttaagctgctg |
| R3.2.4 | ccactgtgtgtgactaagggAACGCGCCTTACCGGATACCT |
| R3.2.5 | ccgaagttggcgactgtaacTTTGAAG |
| R3.2.6 | AATTATATCGCACGcatctgacgaattgacacca |
| R3.2.7 | cagcaggctcaagggtgtatAGAGCACATCCTCATAACGGA |
| R3.2.8 | TTCGTAATCTCGTCGCTGGCAGCCTCCGGCCacttccgattga |
| R3.2.9 | TTCTAAGACAACATTTGAGattcgtcagatggtggcgat |
| R3.2.10 | tatccgccgttacagtcgccGATCCCCGGGTACCGAGCTCACA |
| R3.2.11 | GGTCATACCGGTTTTTTTTTTCTGCCAGCACtaacactaaccgcatatcat |
| R3.2.12 | attaactccccttagtcacaGCGTGCCTG |
| R3Q.2.1 | GTCTACATACTTCCGATTGAATGAGAAAGCGC |
| R3Q.2.2 | GCGTGAGCTTCAAGGGTGTATCTGCTATT |
| R3Q.2.3 | CAGCAGCTTAACACTAACCGGGGAGACAGGGTTCTGAGGTT |
| R3Q.2.4 | AGGTATCCGGTAAGGCGCGTTCCCTTAGTCACACACAGTGG |
| R3Q.2.5 | CTTCAAAGTTACAGTCGCCAACTTCGG |
| R3Q.2.6 | TGGTGTCAATTCGTCAGATGCGTGCGATATAATT |
| R3Q.2.7 | TCCGTTATGAGGATGTGCTCTATACACCCTTGAGCCTGCTG |
| R3Q.2.8 | TCAATCGGAAGTGGCCGGAGGCTGCCAGCGACGAGATTACGAA |
| R3Q.2.9 | ATCGCCACCATCTGACGAATCTCAAATGTTGTCTTAGAA |
| R3Q.2.10 | TGTGAGCTCGGTACCCGGGGATCGGCGACTGTAACGGCGGATA |
| R3Q.2.11 | ATGATATGCGGTTAGTGTTAGTGCTGGCAGAAAAAAAAAACCGGTATGACC |
| R3Q.2.12 | CAGGCACGCTGTGACTAAGGGGAGTTAAT |
| R3.3.1 | tactaaacgtgcACCCTGCCGGCTGAGGTCGT |

| | |
|---|---|
| R3.3.2 | GGTATCAATTTCTCCTGTTCCGatctcgatcaga |
| R3.3.3 | GCGCTTTCTCATggttcacaacta |
| R3.3.4 | ctatttgttaaaAGCTCACGC |
| R3.3.5 | CAAATCAACACGACCTGCTAAcggacaacctgt |
| R3.3.6 | ccttaagcatccAATGTTTCCTGGAAGCTCCCT |
| R3.3.7 | AACCTCAGAACCCTGTCTCCCcggttagtgtta |
| R3.3.8 | tgtgactaagggAACGCGCCTTACCGGATACCT |
| R3.3.9 | cttgttgatcgtTTTGAAG |
| R3.3.10 | AATTATATCGCACGgcaaccttcgtg |
| R3.3.11 | tttaacaaatagACGCGGTCCGTTTTTTCG |
| R3.3.12 | GGGCGACCGATGCTGATTGCCGTTCCGGCAAtagttgtgaacc |
| R3.3.13 | AGAGCACATCCTCATAACGGAtctgatcgagat |
| R3.3.14 | gcacgtttagtaACGTGCCGGACTTGTAGATTCTTCG |
| R3.3.15 | acgatcaacaagGTTTACCAGCGCCAAAATAGAAA |
| R3.3.16 | TCATGGTCATAGcacgaaggttgc |
| R3.3.17 | TTCTAAGACAACATTTGAGtaacactaaccg |
| R3.3.18 | cccttagtcacaGATCCCCGGGTACCGAGCTCACA |
| R3.3.19 | GGTCATACCGGTTTTTTTTTCTGCCAGCACacaggttgtccg |
| R3.3.20 | ggatgcttaaggGCGTGCCTG |
